# Variation in recombination rate and its genetic determinism in sheep (*Ovis Aries*) populations from combining multiple genome-wide datasets

**DOI:** 10.1101/104976

**Authors:** Morgane Petit, Jean-Michel Astruc, Julien Sarry, Laurence Drouilhet, Stéphane Fabre, Carole Moreno, Bertrand Servin

## Abstract

Recombination is a complex biological process that results from a cascade of multiple events during meiosis. Understanding the genetic determinism of recombination can help to understand if and how these events are interacting. To tackle this question, we studied the patterns of recombination in sheep, using multiple approaches and datasets. We constructed male recombination maps in a dairy breed from the south of France (the Lacaune breed) at a fine scale by combining meiotic recombination rates from a large pedigree genotyped with a 50K SNP array and historical recombination rates from a sample of unrelated individuals genotyped with a 600K SNP array. This analysis revealed recombination patterns in sheep similar to other mammals but also genome regions that have likely been affected by directional and diversifying selection. We estimated the average recombination rate of Lacaune sheep at 1.5 cM/Mb, identified about 50,000 crossover hotspots on the genome and found a high correlation between historical and meiotic recombination rate estimates. A genome-wide association study revealed two major loci affecting inter-individual variation in recombination rate in Lacaune, including the *RNF212* and *HEI10* genes and possibly 2 other loci of smaller effects including the *KCNJ15* and *FSHR* genes. Finally, we compared our results to those obtained previously in a distantly related population of domestic sheep, the Soay. This comparison revealed that Soay and Lacaune males have a very similar distribution of recombination along the genome and that the two datasets can be combined to create more precise male meiotic recombination maps in sheep. Despite their similar recombination maps, we show that Soay and Lacaune males exhibit different heritabilities and QTL effects for inter-individual variation in genome-wide recombination rates.

## Introduction

Meiotic recombination is a fundamental biological process that brings a major contribution to the genetic diversity and the evolution of eukaryotic genomes (Baudat and al., 2013). During meiosis, recombination enables chromosomal alignment resulting in proper disjunction and segregation of chromosomes, avoiding deleterious outcomes such as aneuploidy (Hassold, Hall, and Hunt 2007). Over generations, recombination contributes to shaping genetic diversity in a population by creating new allelic combinations and preventing the accumulation of deleterious mutations. Over large evolutionary timescales, divergence in recombination landscapes can lead to speciation: the action of a key actor in the recombination process in many mammals, the gene *PRDM9*, has been shown to have a major contribution to the infertility between two mouse species, making it the only known speciation gene in mammals today (Mihola et al. 2009).

Genetics studies on recombination were first used to infer the organisation of genes along the genome (Sturtevant 1913). With the advance in molecular techniques, more detailed physical maps and eventually whole genome assemblies are now available in many species. The establishment of highly resolutive recombination maps remains of fundamental importance for the validation of the physical ordering of markers, obtained from sequencing experiments (Groenen et al. 2012; Jiang et al. 2014). From an evolutionary perspective the relevant distance between loci is the genetic distance and recombination maps are essential tools for the genetic studies of a species, for estimation of past demography (H. Li and Durbin 2011; Boitard et al. 2016), detection of selection signatures (Sabeti et al. 2002; Voight et al. 2006), QTL mapping (Cox et al. 2009) and imputation of genotypes (Howie, Donnelly, and Marchini 2009) for genome-wide association studies (GWAS) or genomic selection. Precise recombination maps can be estimated using different approaches. Meiotic recombination rates can be estimated from the observation of markers’ segregation in families. Although this is a widespread approach, its resolution is limited by the number of meioses that can be collected within a population and the number of markers that can be genotyped. Consequently highly resolutive meiotic maps have been produced in situations where large segregating families can be studied and genotyped densely (Shifman et al. 2006; Mancera et al. 2008; Groenen et al. 2009; Rockman and Kruglyak 2009; Augustine Kong et al. 2010) or by focusing on specific genomic regions (Cirulli, Kliman, and Noor 2007; Stevison and Noor 2010; Kaur and Rockman 2014). In livestock species, the recent availability of dense genotyping assays has fostered the production of highly resolutive recombination maps (Tortereau et al. 2012; Susan E. Johnston et al. 2016; S. E. Johnston et al. 2017) in particular by exploiting reference population data from genomic selection programs (Sandor et al. 2012a; Ma et al. 2015; Kadri et al. 2016a). Another approach to study the distribution of recombination on a genome is to exploit patterns of correlation between allele frequencies in a population (*i.e.* Linkage Disequilibrium, LD) to infer past (historical) recombination rates (McVean, Awadalla, and Fearnhead 2002; N. Li and Stephens 2003; Chan, Jenkins, and Song 2012). Because the LD-based approach exploits in essence meioses accumulated over many generations, it can provide more precise estimates of local variation in recombination rate. For example, until recently (Pratto et al. 2014; Lange et al. 2016) this was the only indirect known approach allowing to detect fine scale patterns of recombination genome-wide in species with large genomes. Several highly recombining intervals (recombination hotspots) were detected from historical recombination rate maps and confirmed or completed those discovered by sperm-typing experiments (Simon Myers et al. 2005; Crawford et al. 2004). One important caveat of LD-based approaches is that their recombination rate estimates are affected by other evolutionary processes, especially selection that affects LD patterns unevenly across the genome. Hence differences in historical recombination between distant genomic regions have to be interpreted with caution. Despite this, historical and meiotic recombination rates usually exhibit substantial positive correlation (Rockman and Kruglyak 2009; Chan, Jenkins, and Song 2012; Brunschwig et al. 2012; J. Wang et al. 2012).

The LD-based approach does not allow to study individual phenotypes and therefore to identify directly loci influencing inter-individual variation in recombination rates. In contrast, family-based studies in human (A. Kong et al. 2008a; Chowdhury et al. 2009), Drosophila (Stevison and Noor 2010; Chan, Jenkins, and Song 2012) mice (Shifman et al. 2006; Brunschwig et al. 2012) cattle (Sandor et al. 2012a; Ma et al. 2015; Kadri et al. 2016a) and sheep (Susan E. Johnston et al. 2016) have demonstrated that recombination exhibits inter-individual variation and that this variation is partly determined by genetic factors. Two recombination phenotypes have been described: the number of crossovers per meiosis (Genome-wide Recombination Rate, GRR herein) and the fine scale localization of crossovers (Individual Hotspot Usage, IHU herein). GRR has been shown to be influenced by several genes. For example, a recent genome-wide association study found evidence for association with 6 genome regions in cattle (Kadri et al. 2016a). Among them, one of the genomic regions consistently found associated to GRR in mammals is an interval containing the *RNF212* gene. In contrast to GRR, the IHU phenotype seems mostly governed by a single gene in most mammals, *PRDM9*. This zinc-finger protein has a key role in recruiting *SPO11*, thereby directing DNA double-strand breaks (DSBs) that initiate meiotic recombination. Because *PRDM9* recognizes a specific DNA motif, the crossover events happen in hotspots carrying this motif. This *PRDM9* associated process is however not universal, as it is only active in some mammals; canids for example do not carry a functional copy of *PRDM9* and exhibit different patterns for the localization of recombination hotspots (Auton et al. 2013).

As mentioned above, recombination was studied recently in sheep (Susan E. Johnston et al. 2016), which lead to the production of precise genome-wide recombination maps, revealed a similar genetic architecture of recombination rates in sheep as in other mammals and identified two major loci affecting individual variation. Quite interestingly, one the QTL identified in this study, localized near the *RNF212* gene, was clearly demonstrated to have a sex specific effect. This study was performed in a feral population of sheep which is quite distantly related to continental populations (Kijas et al. 2012) and has not managed by humans for a long time. To understand how recombination patterns and genetic determinism can vary across populations, we conducted in this work a study in another sheep population, the Lacaune, from south of France. The Lacaune breed is the main dairy sheep population in France, its milk being mainly used for the production of Roquefort cheese. Starting in 2011, a large genotyping effort started in the breed to implement a genomic selection program (Baloche et al. 2014), and young selection candidates are now routinely genotyped for a medium density genotyping array (about 50K SNP). This constitutes a large dataset of genotyped families that can be used to study recombination, although limited to one sex as only males were used for genomic selection in this population. This dataset offers an opportunity to study variation in recombination and its genetic determinism between very diverged populations of the same species. Hence, a first objective of this study was to elucidate whether these two sheep populations had similar distribution of recombination on the genome and whether they shared the same genetic architecture of the trait, and in particular the same QTLs effects.

The second objective of this study was to compare different approaches to study recombination from independent data in the same population. To this end, in addition to the pedigree data, we exploit a sample of 51 unrelated individuals genotyped with a high-density genotyping array (about 500K SNP). While, the family data was used to establish meiotic recombination maps, the sample of densely genotyped individuals was used to create historical recombination maps of higher resolution. This offered the opportunity to evaluate to which extent sheep ancestral recombination patterns match contemporary ones.

## Materials and Methods

### Study Population and Genotype Data

In this work, we exploited two different datasets of sheep from the Lacaune breed: a pedigree dataset of 8,085 related animals genotyped with the medium density Illumina Ovine Beadchip^®^ including 54,241 SNPs, and a diversity dataset of 70 unrelated Lacaune individuals selected as to represent population genetic diversity, genotyped with the high density Illumina Ovine Infinium^®^ HD SNP Beadchip including (Rochus et al. 2017; Moreno-Romieux et al. 2017)

Standard data cleaning procedures were carried out on the pedigree dataset using plink 1.9 (Chang et al. 2015), excluding animals with call rates below 95% and SNPs with call freq below 98%. After quality controls we exploited genotypes at 46,813 SNPs and 5,940 meioses. For these animals, we only selected the sires which had their own sire known and at least 2 offspring and the sires which did not have their own sire known, but at least 4 offspring. Eventually, 345 male parents, called focal individuals (FIDs) hereafter, met these criteria: 210 FIDs had their father genotype known while the remaining 135 did not (Figure 1).

**Figure 1.**
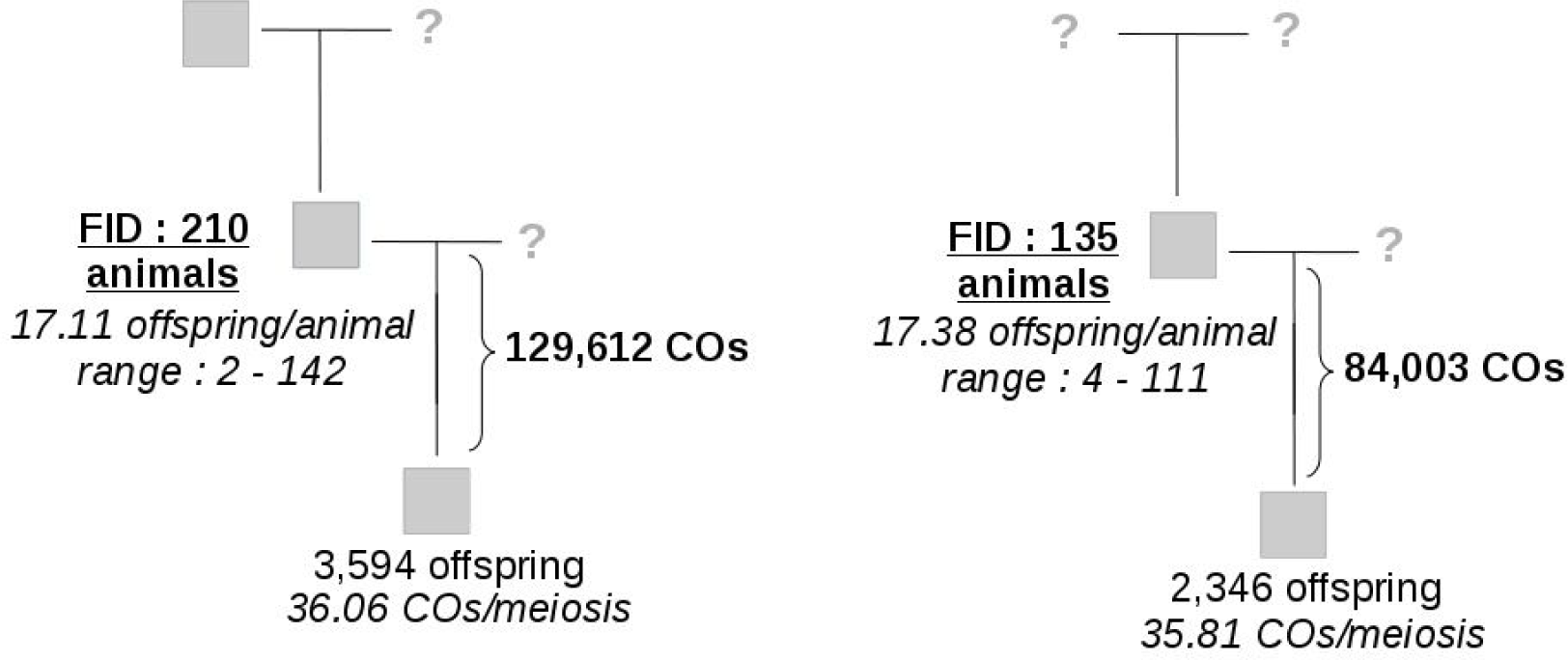
Families used to infer crossovers events. Crossover events were identified in meioses of 345 focal individuals (FIDs). 210 FID had their father known (left) while 135 FID did not (right).

## Recombination Maps

### Meiotic recombination maps from pedigree data

#### Detection of crossovers

Crossover locations were detected using LINKPHASE (Druet and Georges 2015). From the LINKPHASE outputs (*recombination_hmm* files), we extracted crossovers boundaries. We then identified crossovers occurring in the same meiosis less than 3 Mb apart from each other (that we call double crossovers) and considered them as dubious. This number was chosen as it corresponded to clear outliers in the distribution of inter-crossover distances. They are also quite unlikely under crossover interference. We applied the following procedure: given a pair of double crossovers, we set the genotype of the corresponding offspring as missing in the region spanned by the most extreme boundaries and re-run the LINKPHASE analysis. After this quality control step, we used the final set of crossovers identified by LINKPHASE to estimate recombination rates. This dataset consisted of 213,615 crossovers in 5,940 meioses.

#### Estimation of recombination rates

Based on the inferred crossover locations, meiotic recombination rates were estimated in windows of one megabase and between marker intervals of the medium SNP array using the following statistical model, inspired by (Cheung et al. 2007). For small genetic intervals such as considered here, the recombination rate (termed c in the following), is usually expressed in centiMorgans per megabase and the probability that a crossover occurs in one meiosis in an interval *j* (measured in Morgans) is 0.01c_j_l_j_ where l_j_ is the length of the interval expressed in megabases. When considering M meioses, the expected number of crossovers in the interval is 0.01c_j_l_j_M. When combining observations in multiple individuals, we want to account for the fact that they have different average numbers of crossovers per meiosis (termed R_s_ for individual *s*). To do so we multiply the expected number of crossovers in the interval by an individual specific factor equal to (R_s_/R) where R is the average number of crossovers per meiosis among all individuals. Finally, for individual *s* in interval *j* the expected number of crossovers is 0.01c_j_l_j_M_s_R_s_/R. Given this expected number, a natural distribution to model the number of crossovers observed in an interval is the Poisson distribution so that the number y_sj_ of crossovers observed in the interval *j* for an individual *s* is modelled as:

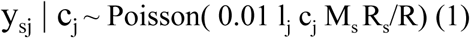

To combine crossovers across individuals, the likelihood for *c_j_* is the product of poisson likelihoods from equation (1).

We then specify a prior distribution for *c_j_*:

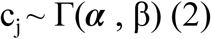

To set α and ß, first raw *c_j_* estimates are computed using the method of (Sandor et al. 2012a) across the genome and then a gamma distribution is fitted to the resulting genome-wide distribution (Figure S1). Combining the prior (2) with the likelihoods in equation (1), the posterior distribution for *c_j_* is:

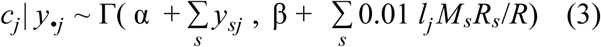

As the localization of crossovers was usually not good enough to assign them with certainty to a single genomic interval, final estimates of *c_j_* are obtained as follows:

i. for each crossover overlapping interval *j* and localized within a window of size *L*, let *x_c_* be an indicator variable that takes value 1 if the crossover occurred in interval *j* and 0 otherwise. Assuming that, locally, recombination rate is proportional to physical distance, set *P*(*x_c_* = 1) = *min*(*l_j_*/*L*, 1).
ii. Using the probability in step (i), sample *x_c_* for each crossover overlapping interval j and set 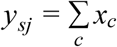
iii. Given *y_sj_*, sample *c_j_* from equation (3)

For each interval considered, perform step (ii) and (iii) above 1000 times to draw samples from the posterior distribution of *c_j_* thereby accounting for uncertainty in the localization of crossovers.

### Historical recombination maps from the diversity data

The diversity data contains 70 Lacaune individuals genotyped for a High Density SNP array comprising 527,823 autosomal markers (Rochus et al. 2017). Nineteen of these individuals are FIDs in the pedigree data. To perform the LD-based analysis on individuals unrelated to the pedigree study, these individuals were therefore removed from the dataset and the subsequent analyses performed on the 51 remaining individuals. Population-scaled recombination rates were estimated using PHASE (N. Li and Stephens 2003). For computational reasons and to allow for varying effective population size along the genome, estimations were carried out in 2 Mb windows, with an additional 100 Kb on each side overlapping with neighbouring windows, to avoid border-effect in the PHASE inference. PHASE was run on each window with default options, except that the number of main iterations was increased to obtain larger posterior samples for recombination rate estimation (option −X10) as recommended in the documentation.

From the PHASE output, 1000 samples were obtained from the posterior distribution of:

- The background recombination rate: ρ_w_ = 4N_w_c_w_, where N_w_ is the effective population size in the window, c_w_ is the recombination rate comparable to the family-based estimate.
- An interval specific recombination intensity λ_j_, for each marker interval *j* of length *l_j_* in the window, such that the population scaled genetic length of an interval is: δ*_j_* = ρ_*w*_λ_*j*_*l_j_*

The medians were used as point estimates of parameters and, computed over the posterior λ_*j*_ and δ_*j*_, computed over the posterior distributions 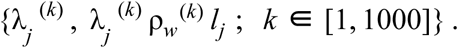

### Identification of intervals harbouring crossover hotspots

Intervals that showed an outlying λ_j_ value compared to the genome-wide distribution of λ_j_ were considered as harbouring a crossover hotspot. Specifically, a mixture of Gaussian distribution was fitted to the genome-wide distribution of *log*_10_ (λ_*j*_) using the mclust R package (Fraley and Raftery 2002), considering that the major component of the mixture modelled the background distribution of λ_j_ in non-hotspots intervals. From this background distribution, a p-value was computed for each interval that corresponded to the null hypothesis that it does not harbour a hotspot. Finally, hotspot harbouring intervals were defined as those for which FDR(λ_j_) < 5%, estimating FDR with the (Storey and Tibshirani 2003) method, implemented in the R qvalue package. This procedure is illustrated in Figure S2.

### Combination of meiotic and historical recombination rates and construction of a high resolution recombination map

To construct a meiotic recombination map of the HD SNP array requires that the historical recombination rate estimates be scaled by 4 times the effective population size. Due to evolutionary pressures, the effective population size varies along the genome, so it must be estimated locally. This can be done by exploiting the meiotic recombination rate inference obtained from the pedigree data analysis as explained below.

Consider a window of one megabase on the genome, using the approach described above, we can sample values c_jk_ (window *j*, sample *k*) from the posterior distribution of the meiotic recombination rate *c_j_*. Similarly, using output from PHASE we can extract samples from the posterior ρ_*jk*_ from the posterior distribution of the historical recombination rates (ρ_*j*_ = ρ_*w*_ λ_*j*_). Now, considering that ρ_*j*_ = 4*Ne_j_c_j_* where *Ne_j_* is the local effective population size of window *j*, we get *log*(ρ_*j*_) = *log*(*Ne_j_*) + *log*(*c_j_*). This justifies using a model on both c_jk_ and ρ_jk_ values:

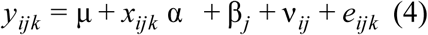

where *y_ijk_* is *log*(*c_jk_*) when i=1 (meiotic-recombination rate sample) and *y_ijk_* is *log*(ρ_*jk*_) when i=2 (historical recombination rate sample). In this model, μ estimates the log of the genome-wide recombination rate, x_ijk_=1 if i=2 and 0 otherwise so that α estimates log(4Ne), where Ne is the average effective population size of the Lacaune population, μ + β_*j*_ estimates log(c_j_) combining population and meiotic recombination rates, and α+(ν_2j_-ν_1j_) estimates log(4Ne_j_). μ and were α_*i*_ were considered as fixed effects while β_*j*_ and ν_*ij*_ and were considered as independent random effects. Using this approach allows to combine in a single model LD- and pedigree-based inferences, while accounting for their respective uncertainties as we exploit posterior distribution samples.

Model (4) was fitted on 20 samples of the posterior distributions of *c_j_* and ρ_*j*_ for all windows of one megabase covering the genome, with an additional fixed effect for each chromosome, using the lme4 R package (Bates et al. 2015). Windows lying less than 4 Mb from each chromosome end were not used because inference on *c_j_* was possibly biased in these regions (see Results). After estimating this model, historical recombination rate estimates of HD intervals were scaled within each window by dividing them by their estimated local effective population size (*i.e.* 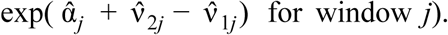 For windows lying within 4 Mb of the chromosome ends, historical recombination rate estimates were scaled using the genome-wide average effective population size 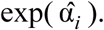 This led eventually to estimates of the meiotic recombination rates, expressed in centiMorgans per megabase, for all intervals of the HD SNP array, which we termed a high resolution recombination map.

#### Effect of recombination hotspots on the recombination rate

For each interval of the medium density SNP array, we computed the number of significant hotspots detected as explained above and the hotspot density (number of hotspots per unit of physical distance). After having corrected for the chromosome effect, the GC content effect and for windows farther than 4 Mb of the chromosome end, we fitted a linear regression model to estimate the effect of hotspots density on the meiotic recombination rate.

### Comparison with Soay sheep recombination maps and integration of the two datasets to produce new male recombination maps in Sheep

In order to compare the recombination maps in Lacaune with the previously established maps in Soay sheep (Susan E. Johnston et al. 2016), we downloaded the raw data from the dryad data repository (doi: 10.5061/dryad.pf4b7) and the additional information available on https://github.com/susjoh/GENETICS_2015_185553. As the approach used in (Susan E. Johnston et al. 2016) to establish recombination maps differs from the one used here, we chose to apply the method of this study to the Soay data to perform a comparison that would not be affected by difference in methods. As the Lacaune data consist only of male meioses, we also only considered male meioses in the Soay data. The final Soay dataset used consisted of 3,445 individuals among which were 299 male FIDs, defined as in the Lacaune analysis. After detecting crossovers with LINKPHASE, one FID exhibited a very high average number of crossover per meiosis (> 100) and was not considered in the analyses (Soay individual ID: RE4844), leaving 298 FIDs. The final dataset consisted of 88,683 crossovers in 2,609 male meiosis and was used to estimated meiotic recombination maps using the exact same approach as described above, both on intervals of one megabase and on the same intervals as the ones considered in the Lacaune meiotic maps on the medium density SNP array. Note that the Soay sheep are not necessarily polymorphic for the same markers as the Lacaune, but that our method is flexible and can nonetheless estimate recombination rates in intervals bordered by monomorphic markers: in such a case adjacent intervals will have the same estimated recombination rate. As the two populations were found to have very similar meiotic recombination maps (see Results), the two sets of crossovers were finally merged to create a combined dataset of 302,298 crossovers in 8,549 male meioses and to estimate new male sheep recombination maps, again on one megabase intervals and on intervals of the medium density SNP array.

### Genome-Wide Association Study on Recombination Phenotypes

#### Genome-wide Recombination Rate (GRR)

The set of crossovers detected was used to estimate the genome-wide recombination rate (GRR) of each FID in the family dataset from their observed number of crossovers per meiosis, adjusting for covariates: year of birth of the parent, considered as a cofactor with 14 levels for years spanning from 1997 to 2010 and insemination month of the offspring’s ewe, treated as a cofactor with 7 levels for months spanning from February to August. We used a mixed-model for estimating the population average GRR μ, covariates fixed effects β and individual breeding values u_s_, while controlling for non genetic individual specific effects a_s_:

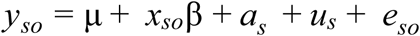

with 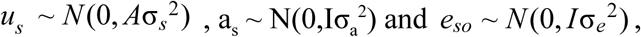 where *y_so_* is the number of crossovers in the meiosis between FID *s* and offspring *o*, A is the pedigree-based relationship matrix between FIDs and *x_so_* the line of the corresponding design matrix for observation *y_so_*. We fitted this model using BLUPf90 (Misztal et al., 2002) and extracted: (i) estimates of variance components σ_*e*_^2^, σ_a_^2^ and σ_*s*_^2^, which allows to estimate the heritability of the trait (calculated as σ_*s*_^2^/(σ_*s*_^2^ + σ_a_^2^ + σ_*e*_^2^)) and (ii) prediction *ũ*_*s*_ of GRR deviation for each FID.

#### Genotype Imputation

Nineteen of the 345 FIDs are present in the diversity dataset of HD genotypes. For the 336 remaining FIDs, their HD genotypes at 507,784 SNPs were imputed with BimBam (Guan and Stephens 2008; Servin and Stephens 2007) using the 70 unrelated Lacaune individuals as a panel. To impute, BimBam uses the fastPHASE model (Scheet and Stephens 2006), which relies on methods using cluster of haplotypes to estimate missing genotypes and reconstruct haplotypes from unphased SNPs of unrelated animal. BimBam was run with 10 expectation-maximization (EM) starts, each EM was run 20 steps on panel data alone, and an additional 1 step on cohort data, with a number of clusters of 15. After imputation BimBam estimates for each SNP in each individual an average number of alleles, termed *mean genotype*, computed from the posterior distribution of the three possible genotypes. This mean genotype has been shown to be efficient for performing association tests (Guan and Stephens 2008). In subsequent analyses, we used the mean genotypes provided by BimBam of the 345 FIDs at all markers of the HD SNP array. To assess the quality of genotype imputation at the most associated regions, 10 markers of the HD SNP array, 1 in chromosome 6 associated region and 9 in the chromosome 7 associated region (see Results) were genotyped for 266 FIDs for which DNA samples were still available. We evaluated the quality of imputation for the most significant SNPs by comparing for each possible genotype its posterior probability estimated by Bimbam to the error rate implied by calling it. We observed a very good agreement between the two measures (Figure S3), which denoted good calibration of the imputed genotypes at top GWAS hits.

#### Single- and multi-QTLs GWAS on GRR

We first tested association of individual breeding values ũ_s_ with mean genotypes at 503,784 single SNPs imputed with BimBam. We tested these associations using the univariate mixed-model approach implemented in the Genome-wide Efficient Mixed Model Association (Gemma) software (Zhou and Stephens, 2012). To accounts for polygenic effects on the trait, the centered genomic relationship matrix calculated from the mean genotypes was used. The p-values reported in the results correspond to the Wald test.

To go beyond single SNP association tests, we also estimated a Bayesian sparse linear mixed-model (Zhou, Carbonetto, and Stephens 2013) as implemented in Gemma. This method allows to consider multiple QTLs in the model, together with polygenic effects at all SNPs. The principle of the method is to have for each SNP *l* an indicator variable 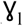 that takes value 1 if the SNP is a QTL and 0 otherwise. The strength of evidence that a SNP is a QTL is measured by the posterior probability 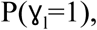, called posterior inclusion probability (PIP). Note that all SNPs are included in the model when doing so. Inference of the model parameters is performed using an iterative MCMC algorithm: the number of iterations was set to 10 millions and inference was made on samples extracted every 100 iterations. When a genome region harbors a QTL, multiple SNP in the region can have elevated PIPs. To summarize the strength of evidence for a *region* to carry a QTL, we calculated a rolling sum of PIPs over 50 consecutive SNPs using the rollsum function of the R zoo package (Zeileis and Grothendieck 2005). Given that the average physical distance between SNPs on the high-density SNP array is about 5 kilobases, this procedure interrogates the probability of the presence of a QTL in overlapping windows of approximately 250 kilobases.

For the univariate analysis, the False Discovery Rate was estimated using the ash package (Stephens 2017) and SNPs corresponding to an FDR < 10% were deemed significant and annotated. For the multivariate analysis, regions where the rolling sum of PIPs exceeded 0.15 were further annotated. The annotation of the QTL regions consisted in extracting all genes from the Ensembl annotation v87 along with their Gene Ontology (GO) annotations and interrogated for their possible involvement in recombination.

### Variant Discovery and Additional Genotyping in *RNF212*

#### Identification and assignation of the *RNF212* sheep genome sequence

The *RNF212* gene was not annotated on the *Ovis aries* v3.1 reference genome. Nevertheless, a full sequence of *RNF212* was found in the scaffold01089 of *Ovis orientalis* (assembly Oori1, NCBI accession NW_011943327). By BLAST alignment of this scaffold, ovine *RNF212* could be located with confidence on chromosome 6 in the interval OAR6:116426000-116448000 of Oari3.1 reference genome (Figure S4). This location was confirmed by BLAST alignment with the bovine *RNF212* gene sequence. We also discovered that the Oari3.1 unplaced scaffold005259 (NCBI accession JH922970) contained the central part of *RNF212* (exons 4-9) and it could be placed within a large assembly gap. Moreover, we also observed that automatically annotated non-coding RNA in the *RNF212* interval matched exonic sequence of *RNF212* (Figure S4).

#### Variant discovery in *RNF212* in the Lacaune population

Based on the genomic sequence and structure of the *RNF212* gene annotated in *Ovis orientalis* (NCBI accession NW_011943327), a large set of primers were designed using PRIMER3 software (Table S1) for amplification of each annotated exon and some intron part corresponding to exonic region annotated in *Capra hircus* (Chir_v1.0). PCR amplification (GoTaq, Promega) with each primer pair was realized on 50ng of genomic DNA from 4 selected homozygous Lacaune animals exhibiting the GG and AA (non imputed) genotypes at the most significant SNP of the medium density SNP array of the chromosome 6 QTL (rs418933055, p-value 2.56e-17). Each PCR product was sequenced via the BigDye Terminator v3.1 Cycle Sequencing kit and analyzed on an ABI3730 sequencing machine (Applied Biosystems). Sequenced reads were aligned against the *Ovis orientalis RNF212* gene using CLC Main Workbench Version 7.6.4 (Qiagen Aarhus) in order to identify polymorphisms.

#### Genotyping of mutations in *RNF212*

The genotyping of 266 genomic DNA from Lacaune animals for the four identified polymorphisms within the ovine *RNF212* gene was done by Restriction Fragment Length Polymorphism (RFLP) after PCR amplification using dedicated primers (Table S1) (GoTaq, Promega), restriction enzyme digestion (BsrBI for SNP_14431_AG; RsaI for SNP_18411_GA; and Bsu36I for both SNP_22570_CG and SNP_22594_AG; New England Biolabs) and resolution on 2% agarose gel.

## Results

### High-Resolution Recombination Maps

#### Meiotic recombination maps: genome-wide recombination patterns

We studied meiotic recombination using a pedigree of 6,230 individuals, genotyped for a medium density SNP array (50K) comprising around 54,000 markers. After quality controls we exploited genotypes at 46,813 SNPs and identified 213,615 crossovers in 5,940 meioses divided among 345 male parents (FIDs) (see Methods). The pedigree information available varied among focal individuals (Figure 1): 210 FIDs had their father genotype known while the remaining 135 did not. Having a missing parent genotype did not affect the detection of crossovers as the average number of crossovers per meiosis in the two groups was similar (36.1 with known father genotype and 35.8 otherwise) and the statistical effect of the number of offspring on the average number of crossovers per meiosis was not significant (p>0.23). This can be explained by the fact that individuals that lacked father genotype information typically had a large number of offspring (17.4 on average, ranging from 4 to 111), allowing to infer correctly their haplotype phase from their offspring genotypes only. Overall, given that the physical genome size covered by the medium density SNP array is 2.45 gigabases, we estimate that the mean recombination rate in our population is about 1.5 cM/megabase.

Based on the crossovers identified, we developed a statistical model to estimate meiotic recombination rates (see methods) and constructed meiotic recombination maps at two different scales: for windows of one megabase and for each interval of the medium density SNP array. As this statistical approach allowed to evaluate the uncertainty in recombination rate estimates, we provide respectively in File S1 and S2, along with the recombination rate estimates in each interval, their posterior variance and 90% credible intervals. Graphical representation of the meiotic recombination maps of all autosomes are given in File S3.

The recombination rate on a particular chromosome region was found to depend highly on its position relative to the telomere and to the centromere for metacentric chromosomes, *i.e.* chromosomes 1, 2 and 3 in sheep (Figure S5). Specifically, for acrocentric and metacentric chromosomes, recombination rate estimates were elevated near telomeres and centromeres, but very low within centromeres. In our analysis, recombination rate estimates were found low in intervals lying within 4 megabases of chromosome ends. While this could represent genuine reduction in recombination rates near chromosome ends it is also likely due to crossovers being undetected in our analysis as only few markers are informative to detect crossovers at chromosome ends. In the following analyses, we therefore did not consider regions lying within 4 Mb of the chromosomes ends.

From local recombination rate estimates in 1 Mb windows or medium SNP array intervals, we estimated chromosome specific recombination rates (Figure S6). Difference in recombination rates between chromosomes was relatively well explained by their physical size, larger chromosomes exhibiting smaller recombination rates. Even after accounting for their sizes, some chromosomes showed particularly low (chromosomes 9, 10 and 20) or particularly high (chromosomes 11 and 14) recombination rates. In low recombining chromosomes, large regions had very low recombination, between 9 and 14 Mb on chromosome 9, 36 and 46 Mb on chromosome 10 and between 27 and 31 Mb on chromosome 20. In highly recombining chromosomes, recombination rates were globally higher on chromosome 14, while chromosome 11 exhibited two very high recombination windows between 7 Mb and 8 Mb and between 53 and 54 Mb. In addition, we found, consistent with the literature, that GC content was quite significantly positively correlated with recombination rate both in medium SNP array intervals (p-value < 10 ^-16^, r=0.20) and in 1 Mb intervals (p-value < 10 ^-16^, r=0.28).

#### Estimation of historical recombination rates and identification of crossover hotspots

We used a different dataset, with 51 unrelated individuals from the same Lacaune population genotyped for the Illumina HD SNP array (600K) comprising 527,823 autosomal SNPs after quality controls. Using a multipoint model for LD patterns (N. Li and Stephens 2003), we estimated, for each marker interval of the HD SNP array, historical recombination rates (see Methods). Compared to meiotic maps, these estimates offer a greater precision as they in essence exploit meioses cumulated over many generations. However, the historical recombination rates obtained are scaled by the effective population size (ρ = 4*N_2_c* where Ne is the effective population size and c the meiotic recombination rate) which is unknown, and may vary along the genome due to evolutionary pressures, especially selection. Thanks to the higher precision in estimation of recombination rate, LD-based recombination maps offer the opportunity to detect genome intervals likely to harbour crossover hotspots. A statistical analysis of historical recombination rates (see Methods) identified about 50,000 intervals exhibiting elevated recombination intensities (Figure S2) as recombination hotspots, corresponding to an FDR of 5%. From our historical recombination map, we could conclude that 80% crossover events occurred in 40% of the genome and that 60% of crossover events occurred in only 20% of the genome (Figure S7).

#### High-resolution recombination maps combining family and population data

Having constructed recombination maps with two independent approaches and having datasets in the same population of Lacaune sheep allowed first to evaluate to which extent historical crossover hotspots explain meiotic recombination, and second to estimate the impact of evolutionary pressures on the historical recombination landscape of the Lacaune population. We present our results on these questions in turn.

We studied whether variation in meiotic recombination can be attributed to the historical crossover hotspots detected from LD patterns only. For each interval between two adjacent SNPs of the medium density array, we (i) extracted the number of significant historical hotspots and (ii) calculated the historical hotspot density (in number of hotspots per unit of physical distance). We found both covariates to be highly associated with meiotic recombination rate estimated on family data (r=0.15 with hotspot density (p < 10 ^-16^) and r=0.19 with the number of hotspots (p<10 ^-16^)). These correlations hold after correcting for chromosome and GC content effects (respectively r=0.14 (p<10 ^-16^) and r=0.18 (p<10 ^-16^)). Figure 2 illustrates this finding in two one-megabase intervals from chromosome 24, one that exhibits a very high recombination rate (7.08 cM/Mb) and the second a low one (0.46 cM/Mb). In this comparison, the highly recombining window carries 36 recombination hotspots while the low recombinant one exhibits none. As the historical background recombination rates in the two windows are similar (0.7/Kb for the one with a high recombination rate, and 0.2/Kb for the other), the difference in recombination rate between these two regions is largely due to their contrasted number of historical crossover hotspots.

**Figure 2.**
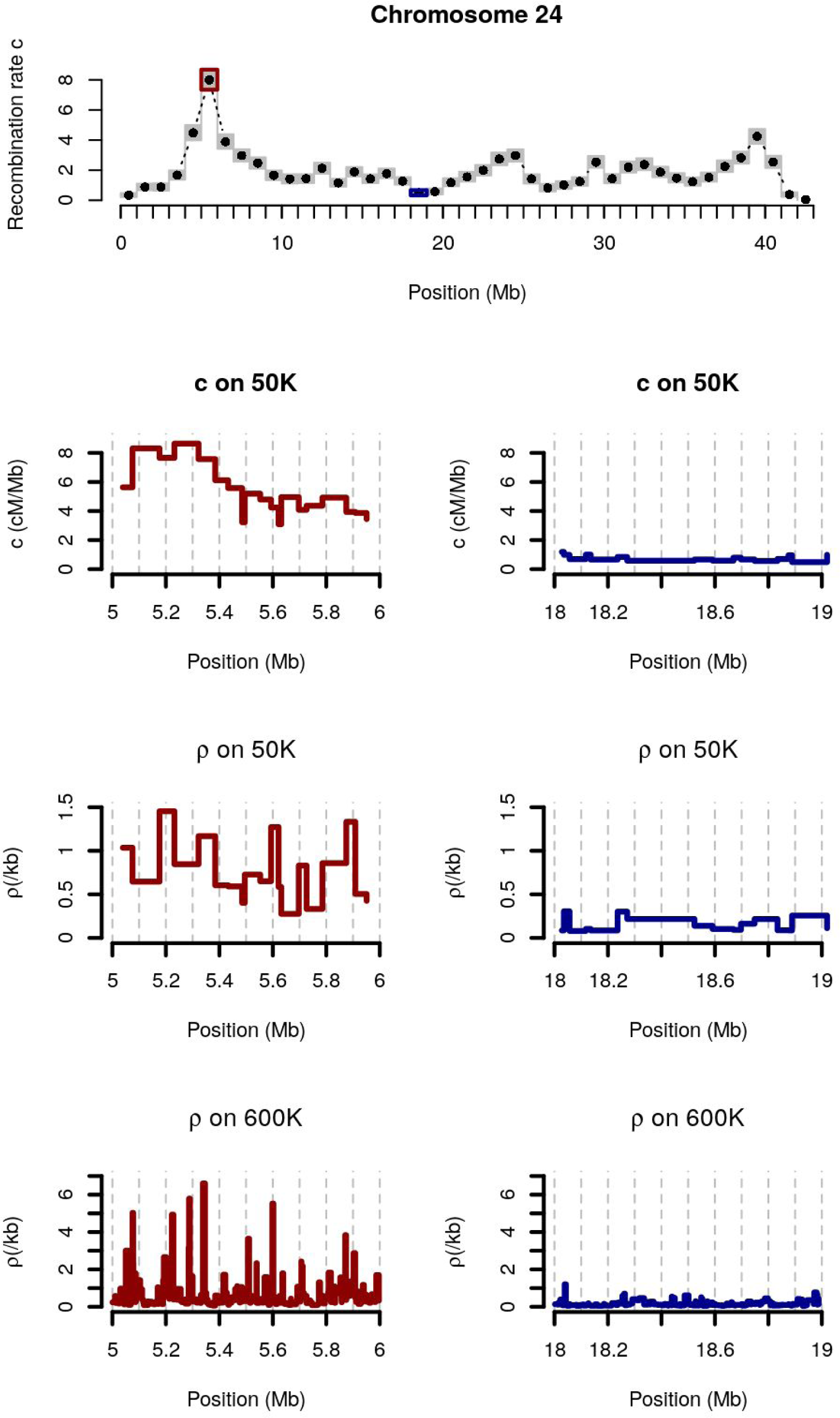
Comparison between population-based recombination rate and meiotic recombination rate for two 1 Mb windows on Sheep chromosome 24. Top: meiotic recombination rate along chromosome 24. Two windows with high (left,red) and low (right,blue) meiotic recombination rates estimates are zoomed in. Each panel represents, from top to bottom: meiotic recombination rate estimates (c) in SNP intervals of the 50K SNP array, population-based recombination rate estimates (ρ) in SNP intervals of the 50K SNP array and population-based recombination rate estimates (ρ) in SNP intervals of the HD (∼600K) SNP array.

In order to study more precisely the relationship between historical and meiotic recombination rates, we fitted a linear mixed model (see Methods) that allowed to estimate the average effective population size of the population, the correlation between meiotic and historical recombination rates and to identify genome regions where historical and meiotic recombination rates were significantly different. We found the effective population size of the Lacaune population to be about 7,000 individuals and a correlation of 0.73 between meiotic and historical recombination rates (Figure 3). We discovered 7 regions where historical recombination rates were much lower than meiotic ones and 3 regions where they were much higher (Table 1, Figure S8). Seven of these 10 regions have extreme recombination rates compared to other genomic regions. To quantify to which extent a window is extreme, we indicate in Table 1, for each window, the proportion of the genome with a lower recombination rate (q_w_). For 6 of these 7 regions, the historical recombination rate is more extreme than the meiotic rate: four regions have very low meiotic recombination rate and even lower historical recombination rates (the two regions on chromosome 3 and two regions on chromosome 10, between 36-37 megabases and between 42-44 megabases); two regions have very high meiotic recombinations rates and even higher historical recombination rates (on chromosome 12 and on chromosome 23). For these six regions, the discrepancy between meiotic recombination and historical recombination estimates can be explained by the fact that we used a genome-wide prior in our model to estimate meiotic recombination rates that has the effect of shrinking our estimates toward the mean. Because historical estimates were not shrunk in the same way, for these six outlying regions the two estimates did not concur and it is possible that our meiotic recombination rate estimates were slightly over (resp. under) estimated. Out of the four remaining outlying windows, three had a low historical recombination rate but did not have particularly extreme meiotic recombination rates, so that the effect of shrinkage is not likely to explain the discrepancy between meiotic and historical recombination rates. Indeed, these three regions corresponded to previously identified selection signatures in sheep: a region on chromosome 6 spanning 2 intervals between 36 and 38 megabases contains the *ABCG2* gene, associated to milk production (Cohen-Zinder et al. 2005), and the *LCORL* gene associated to stature (recently reviewed in (Takasuga 2015)). This region has been shown to have been selected in the Lacaune breed (Fariello et al. 2014; Rochus et al. 2017); a region spanning one interval on chromosome 10, between 29 and 30 megabases contains the *RXFP2* gene, associated to polledness and horn phenotypes (Susan E. Johnston et al. 2013) and found to be under selection in many sheep breeds (Fariello et al. 2014); and a region on chromosome 13 between 63 and 64 megabases that contains the *ASIP* gene responsible for coat color phenotypes in many breeds of sheep (Norris and Whan 2008), again previously demonstrated to have been under selection. For these three regions, we explain the low historical recombination estimates by a local reduction of the effective population size due to selection.

**Table 1.** **Genome regions where meiotic and population-based recombination rates differ significantly.** ρ^*s*^: population-based recombination rate, c: meiotic recombination rate. ρ^*s/c*^: ratio of population to meiotic recombination rate. q_w_: proportion of genome regions with lower meiotic recombination rate. Details on the estimation of these parameters are given in the text. Regions with p-values ≤ 10^-4^ were considered outliers (FDR = 0.02). Regions in bold correspond to potential selection signatures.

| Chromosome | Window span (Mb) | $q_w$ | p-value | $\rho^s/c$ |
| --- | --- | --- | --- | --- |
| 3 | 103-104 | 0.06 | $1.6 \cdot 10^{-5}$ | 0.28 |
| 3 | 109-110 | 0.04 | $1.8 \cdot 10^{-5}$ | 0.28 |
| **6** | **36-38** | **0.14** | **$1.2 \cdot 10^{-7}$** | **0.22** |
| **10** | **29-30** | **0.77** | **$8.8 \cdot 10^{-5}$** | **0.31** |
| 10 | 36-37 | 0.01 | $2.1 \cdot 10^{-5}$ | 0.29 |
| 10 | 42-44 | <0.01 | $1.2 \cdot 10^{-14}$ | 0.11 |
| **13** | **63-64** | **0.33** | **$7.4 \cdot 10^{-6}$** | **0.31** |
| 12 | 4-5 | 0.92 | $7.4 \cdot 10^{-6}$ | 3.7 |
| **20** | **28-29** | **0.01** | **$1.7 \cdot 10^{-5}$** | **3.6** |
| 23 | 10-11 | 0.97 | $5.1 \cdot 10^{-6}$ | 3.8 |

**Figure 3.**
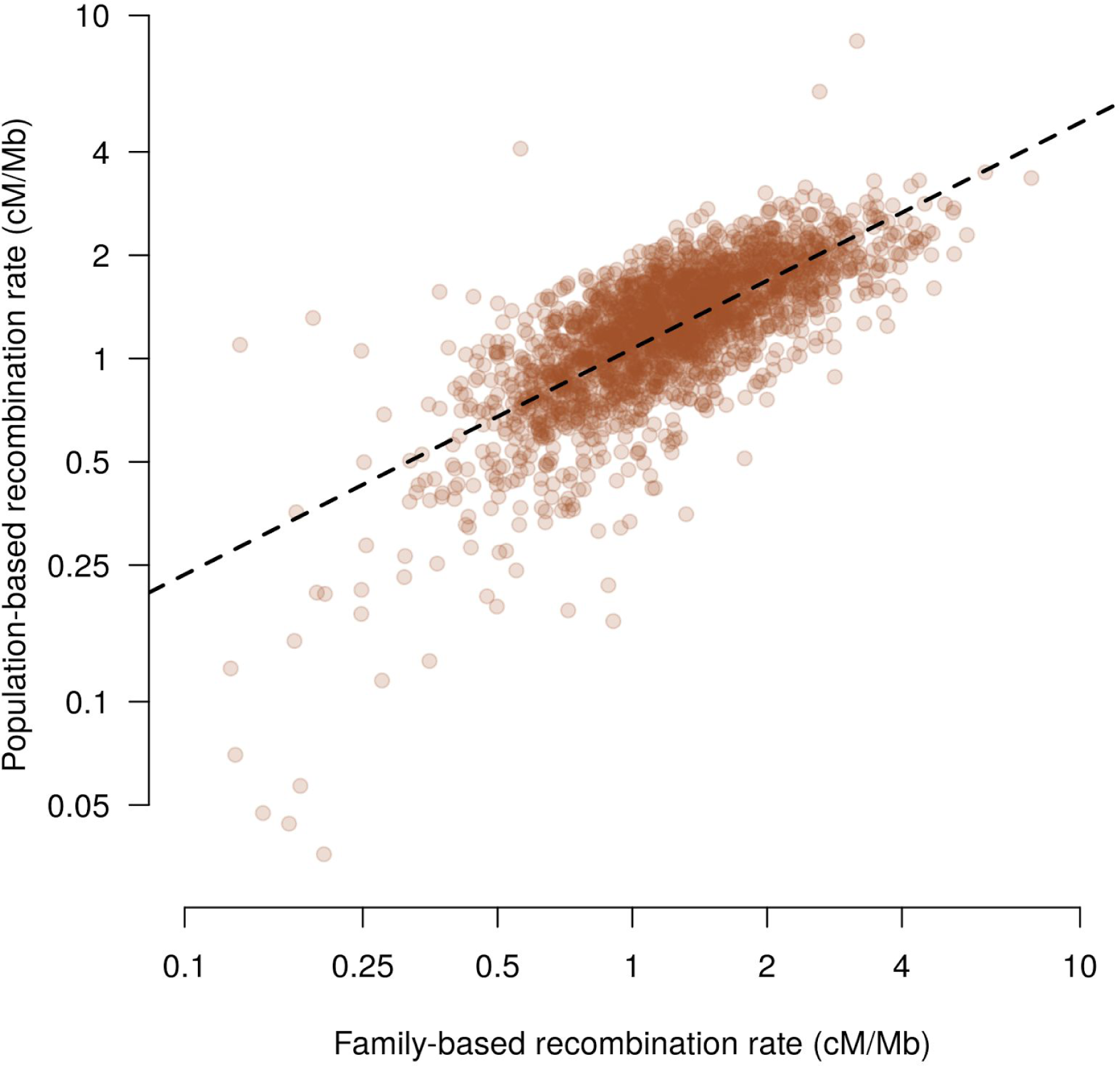
Population-based and meiotic recombination rates in windows of one megabase. The dashed line is the regression for population recombination rate on the family recombination rate. Values are shown on a logarithmic scale.

Finally, one of the three regions with a high historical recombination rate, on chromosome 20 between 28 and 29 megabases had a low meiotic recombination rate, so that the effect of shrinkage cannot explain the discrepancy. This region harbours a cluster of olfactory receptors genes and its high historical recombination rate could be explained by selective pressure for increased genetic diversity in these genes (*i.e.* diversifying selection), a phenomenon which has been shown in other species (*e.g.* pig (Groenen et al. 2012), human (Ignatieva et al. 2014), rodents (Stathopoulos, Bishop, and O’Ryan 2014)). Finally, we used the meiotic recombination rates to scale the historical recombination rate estimates and produce high-resolution recombination maps on the HD SNP array (Supporting File S4).

#### Improved male recombination maps by combining Lacaune and Soay sheep data

Recently, recombination maps have been estimated in another sheep population, the Soay (Susan E. Johnston et al. 2016). Soay sheep is a feral population of ancestral domestic sheep living on an island located northwest of Scotland. The Lacaune and Soay populations are genetically very distant, their genome-wide Fst, calculated using the sheephapmap data (Kijas et al. 2012), being about 0.4. Combining our results with results from the Soay offered a rare opportunity to study the evolution of recombination over a relatively short time scale as the two populations can be considered separated at most dating back to domestication, about 10,000 years ago. The methods used in the Soay study are different from those used here, but the two datasets are similar, although the Soay data has fewer male meioses (2,604 vs. 5,940 in the present study). In order to perform a comparison that would not be affected by differences in estimation methods, we ran the method developed for the Lacaune data to estimate recombination maps on the Soay data. As the Soay study showed a clear effect of sex on recombination rates, we estimated recombination maps on male meioses only. Figure 4 presents the comparison of recombination rates between the two populations in marker intervals of the medium density SNP array. The left panel shows that the two populations exhibit very similar recombination rates (r = 0.82, p<10 ^-16^), although Soay recombination rates appear higher for low recombining intervals (c < 1.5 cM/Mb in gray on the figure). We explain this by the shrinkage effect of the prior, that is more pronounced in the Soay as the dataset is smaller: the right panel on Figure 4 shows that the posterior variance of the recombination rates are clearly higher in Soays for low recombining intervals while they are similar for more recombining intervals. Overall, our results on the comparison of the recombination maps in the two populations are consistent with the two populations having the same amplitude and distribution of recombination on the genome. We therefore analyzed the two populations together to create new male recombination maps based on 302,298 crossovers detected in 8,549 meioses (Supporting file S5). Combining the two dataset together lead to a clear reduction in the posterior variance of the recombination rates, i.e. an increase in their precision (Figure S9).

**Figure 4.**
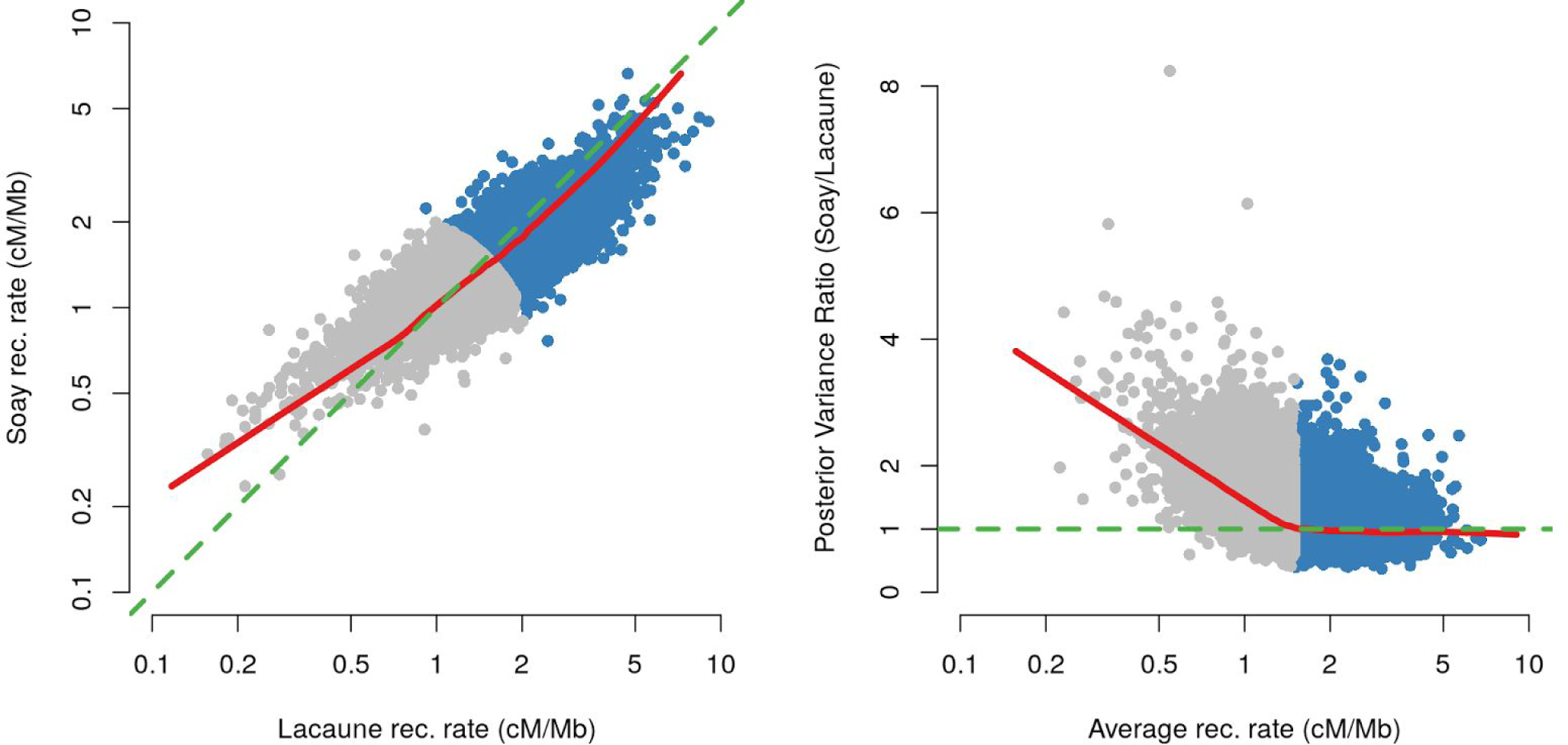
Comparison of recombination rates in Soay and Lacaune populations. Left: scatterplot of posterior means of recombination rates (on a log scale). The green line is the line y=x and the red line is a lowess smoothed line (f=0.05). Right: Scatterplot of the ratio of posterior variance (Soay/Lacaune) as a function of the average of the posterior mean recombination rates in the two populations (on a log scale). The green line corresponds to equal variances and the red line is a lowess smoothed line (f=0.05). Points in gray on both panels correspond to intervals with average recombination rate < 1.5 cM/Mb.

### Genetic Determinism of Genome-wide Recombination Rate in Lacaune sheep

Our dataset provides information on the number of crossovers for a set of 5,940 meioses among 345 male individuals. Therefore, it allows to study the number of crossovers per meiosis (GRR) as a recombination phenotype.

#### Genetic and environmental effects on GRR

We used a linear mixed-model to study the genetic determinism of GRR. The contribution of additive genetic effects was estimated by including a random FID effect with covariance structure proportional to the matrix of kinship coefficients calculated from pedigree records (see Methods). We also included environmental fixed effects in the model: year of birth of the FID and insemination month of the ewe for each meiosis. We did not find significant differences between the FID year of birth, however the insemination month of the ewe was significant (p = 1.3 10 ^-3^). There was a trend in increased recombination rates from February to May followed by a decrease until July and a regain in August although the number of inseminations in August is quite low, leading to a high standard error for this month (Figure S10). Based on the estimated variance components (Table 2), we estimated the heritability of GRR in the Lacaune male population at 0.23.

**Table 2.** **Decomposition of the inter-individual variation in Genome-wide Recombination Rate.**

| **Phenotype** | **Number of sires** | **Genetic Variance** | **Phenotypic Variance** | **Heritability** |
| --- | --- | --- | --- | --- |
| **GRR** | 345 | 6.86<br>(0.75) | 29.73<br>(0.84) | 0.23<br>(0.02) |

Genetic variance, phenotypic variance and heritability of GRR among 345 male individuals. Figures in brackets are standard errors.

#### Genome-wide association study identifies three major loci affecting GRR in Lacaune sheep

The additive genetic values of FIDs, predicted from the above model were used as phenotypes in a genome-wide association study. Among the 345 FIDs with at least two offsprings, the distribution of the phenotype was found to be approximately normally distributed (Figure S11). To test for association of this phenotype with SNPs markers, we used a mixed-model approach correcting for relatedness effects with a genomic relationship matrix (see Methods). Using our panel of 70 unrelated Lacaune, we imputed the 345 FIDs for markers of the HD SNP array. With these imputed genotypes, we performed two analyses. The first was an association test with univariate linear mixed models, which tested the effect of each SNP in turn on the phenotype (results in Supporting File S6) the second fitted a Bayesian sparse linear mixed model, allowing multiple QTLs to be included in the model (results in Supporting File S7).

Figure 4 illustrates the GWAS results: the top plot shows the p-values of the single SNP analysis and the bottom plot, the posterior probability that a region harbours a QTL, calculated on overlapping windows of 20 SNPs. The single SNP analysis revealed six significant regions (FDR < 10 %): two on chromosome 1, one on chromosome 6, one on chromosome 7, one on chromosome 11 and one chromosome 19. Regions of chromosome 6 and 7 exhibited very low p-values whereas the other three showed less intense association signals. The multi-QTLs Bayesian analysis was conclusive for two regions (regions on chromosome 6 and chromosome 7) while the rightmost region on chromosome 1 was suggestive (Table 3). Two additional suggestive regions were discovered on chromosome 3. Using the multi-QTL approach of (Zhou, Carbonetto, and Stephens 2013) allowed to estimate that together, QTLs explain about 40% of the additive genetic variance for GRR, with a 95% credible interval ranging from 28 to 53 %.

**Table 3:** **SNPs associated with GRR.**

| **Rs number** | **Chr** | **Position (bp)** | **Minor allele** | **p** | **$\beta$** | **P-value** | **pQTL** |
| --- | --- | --- | --- | --- | --- | --- | --- |
| **rs430436336** | 1 | 180044043 | A | 0.11 | 2.19 | $8.08 \cdot 10^{-6}$ | 0.006 |
| **rs400472211** | 1 | 268670581 | A | 0.33 | 0.86 | $9.41 \cdot 10^{-6}$ | 0.03 |
| **rs418551122** | 3 | 75216491 | A | 0.30 | 0.76 | $2.42 \cdot 10^{-5}$ | 0.04 |
| **rs407545143** | 3 | 201298545 | G | 0.24 | 1.13 | $9.36 \cdot 10^{-4}$ | 0.07 |
| **rs411987057** | 6 | 116517201 | C | 0.22 | -2.30 | $1.31 \cdot 10^{-16}$ | 0.19 |
| **rs401206888** | 6 | 116440663 | G | 0.14 | -1.95 | $2.04 \cdot 10^{-16}$ | 0.16 |
| **rs412583165** | 6 | 116525709 | G | 0.27 | -2.38 | $9.8 \cdot 10^{-17}$ | 0.15 |
| **rs429477322** | 6 | 116509403 | A | 0.18 | -2.17 | $3.94 \cdot 10^{-16}$ | 0.11 |
| **rs161854895** | 6 | 116491013 | G | 0.22 | -2.17 | $2.53 \cdot 10^{-16}$ | 0.11 |
| **rs398811467** | 6 | 116472870 | A | 0.13 | -1.94 | $2.51 \cdot 10^{-16}$ | 0.14 |
| **rs407110999** | 7 | 22859168 | G | 0.25 | 1.37 | $8.71 \cdot 10^{-7}$ | 0.10 |
| **rs413147562** | 7 | 22798236 | A | 0.23 | 1.61 | $1.20 \cdot 10^{-7}$ | 0.71 |

P-values correspond to the single SNP Wald test. $\beta$ corresponds to the effect of SNP (in number of crossover per meiosis) on GRR and pQTL is the probability for the SNP to be a QTL estimated using a Bayesian Sparse Linear Mixed Model (see Methods).

The most significant region was located on the distal end of chromosome 6 and corresponded to a locus frequently associated to variation in recombination rate. In our study the significant region contained 10 genes: *CTBP1*, *IDUA*, *DGKQ*, *GAK*, *CPLX1*, *UVSSA*, *MFSD7*, *PDE6B*, *PIGG* and *RNF212*. For each of these genes, except *RNF212* which was not annotated on the genome (see below), we extracted their gene ontology of the Ensembl v87 database, but none was clearly annotated as potentially involved in recombination. However, two genes were already reported as having a statistical association with recombination rate: *CPLX1* and *GAK (Augustine Kong et al. 2014)*. *CPLX1* has no known function that can be linked to recombination (Augustine Kong et al. 2014) but *GAK* has been shown to form a complex with the cyclin-G, which could impact recombination (Nagel et al. 2012). However, *RNF212* can be deem a more likely candidate due to its function and given that this gene was associated with recombination rate variation in human (Chowdhury et al. 2009), (A. Kong et al. 2008b), in bovine (Sandor et al. 2012b; Kadri et al. 2016b; Ma et al. 2015) and in mouse (Reynolds et al. 2013). *RNF212* is not annotated in the sheep genome assembly oviAri3, however this chromosome 6 region corresponds to the bovine region that contains *RNF212* (Figure S4). We found an unassigned scaffold (scaffold01089, NCBI accession NW_011943327) of *Ovis orientalis musimon* (assembly Oori1) that contained the full *RNF212* sequence and that could be placed confidently in the QTL region. To confirm *RNF212* as a valid positional candidate, we studied further the association of its polymorphisms with GRR in results presented below.

The second most significant region was located between 22.5 and 23.1 megabases on chromosome 7. All significant SNPs in the region were imputed, *i.e.* the association would not have been found based on association of the medium density array alone. It matched an association signal on GRR in Soay sheep (Susan E. Johnston et al. 2016). Consistent with our finding, in the Soay sheep study, this association was only found using regional heritability mapping and not using single SNP associations with the medium density SNP array. This locus could match previous findings in cattle (association on chromosome 10 at about 20 Mb on assembly btau3.1), however the candidate genes mentioned in this species (*REC8* and *RNF212B*) were located respectively 2 and 1.5 megabases away from our strongest association signal. In addition, none of the SNPs located around these two candidate genes in cattle were significant in our analysis. Eleven genes were present in the region: *OR10G2*, *OR10G3*, *TRAV5*, *TRAV4*, *SALL2*, *METTL3*, *TOX4*, *RAB2B*, *CHD8*, *SUPT16H* and *RPGRIP1*. The study of their gene ontology, extracted from the Ensembl v87 database, revealed that none of them were associated with recombination, although *SUPT16H* could be involved in mitotic DSB repair (Kari et al. 2011). However another functional candidate, *CCNB1IP1*, also named *HEI10*, was located between positions 23,946,971 and 23,951,850 bp, about 500 Kb from our association peak. This gene is a good functional candidate as it has been shown to interact with *RNF212*: *HEI10* allows to eliminate the *RNF212* protein from early recombination sites and to recruit other recombination intermediates involved in crossover maturation (Qiao et al. 2014; Rao et al. 2016). Again SNPs located at the immediate proximity of *HEI10* did not exhibit significant associations with GRR. Hence, our association signal did not allow to pinpoint any clear positional candidate among these functional candidates (see Figure S12). However, it was difficult to rule them out completely for three reasons. First, with only 345 individuals, our study may not be powerful enough to localize QTLs with the required precision. Second, the presence of causal regulatory variants, even at distances of several hundred kilobases is possible. Finally, the associated region of *HEI10* exhibited apparent rearrangements with the human genome, possibly due to assembly problems in oviAri3. These assembly problems could be linked to the presence of genomic sequences coding for the T-cell receptor alpha chain. This genome region is in fact rich in repeated sequences making its assembly challenging. Overall, identifying a single positional and functional candidate gene in this gene-rich misassembled genomic region was not possible based on our data alone.

Our third associated locus was located on chromosome 1 between 268,600 and 268,700 kilobases. In cattle, the homologous region, located at the distal end of cattle chromosome 1, has also been shown to be associated with GRR (Kadri et al. 2016a; Ma et al. 2015). In these studies the *PRDM9* gene has been proposed as a potential candidate gene, especially because it is a strong functional candidate given its proven effect on recombination phenotypes. In sheep, *PRDM9* is located at the extreme end of chromosome 1, around 275 megabases, 7 megabases away from our association signal (Ahlawat et al. 2016). Hence, *PRDM9* was not a good positional candidate for association with GRR in our sheep population. However, the associated region on chromosome 1 contains a single gene, *KCNJ15*, which has been associated with DNA double-strand breaks repair in human cells (Słabicki et al. 2010).

Finally, the two regions on the chromosome 3 were analyzed. The first was located between 75,162 and 75,319 kilobases and contains only one annotated gene coding for the receptor for follicle-stimulating hormone (*FSHR*). Though it does not affect recombination directly it is necessary for the initiation and maintenance of normal spermatogenesis in males (Tapanainen et al. 1997). The second region on the chromosome 3 was located between 201,198 and 201,341 kilobases but does contain any annotated gene.

### Mutations in the *RNF212* gene are strongly associated to Genome-wide Recombination Rate variation in Lacaune sheep

The QTL with the largest effect in our association study corresponded to a locus associated to GRR variation in other species and harbouring the *RNF212* gene. As it was a clear positional and functional candidate gene, we carried out further experiments to interrogate specifically polymorphisms within this gene. As stated above, we used the sequence information available for the *RNF212* gene from *Ovis orientalis* which revealed that *RNF212* spanned 23,7 Kb on the genome and may be composed of 12 exons by homology with bovine *RNF212*. However, mRNA annotation indicated multiple alternative exons. Surprisingly, the genomic structure of ovine *RNF212* was not well conserved with goat, human and mouse syntenic *RNF212* genes (Figure S4). As a first approach, we designed primers for PCR amplification (see Methods) and sequencing of all annotated exons and some intronic regions corresponding to exonic sequences of *Capra hircus RNF212*. By sequencing *RNF212* from 4 carefully chosen Lacaune animals homozygous GG or AA at the most significant SNP of the medium density SNP array on chromosome 6 QTL (rs418933055, p-value 2.56 10 ^-17^), we evidenced 4 polymorphisms within the ovine *RNF212* gene (2 SNPs in intron 9, and 2 SNPs in exon 10). The 4 mutations were genotyped in 266 individuals of our association study. We then tested their association with GRR using the same approach as explained above (results in Supporting File S8) and computed their linkage disequilibrium (genotypic r ^2^) with the most associated SNPs of the high-density genotyping array (see Figure S13) (Table 4). Two of these mutations were found highly associated with GRR, their p-values being of the same order of magnitude (p<10 ^-16^) as the most associated SNP (rs412583165), one of them was even more significant than the most significant imputed SNP (p = 6.25 10 ^-17^ vs p = 9.8 10 ^-17^). We found a clear agreement between the amount of LD between a mutation and the most associated SNPs and their association p-value (see Figure S13). Overall, these results showed that polymorphisms within the *RNF212* gene were strongly associated with GRR, and likely tagged the same causal mutation as the most associated SNP. This confirmed that *RNF212*, a very strong functional candidate, was also a very strong positional candidate gene underlying our association signal.

**Table 4:**
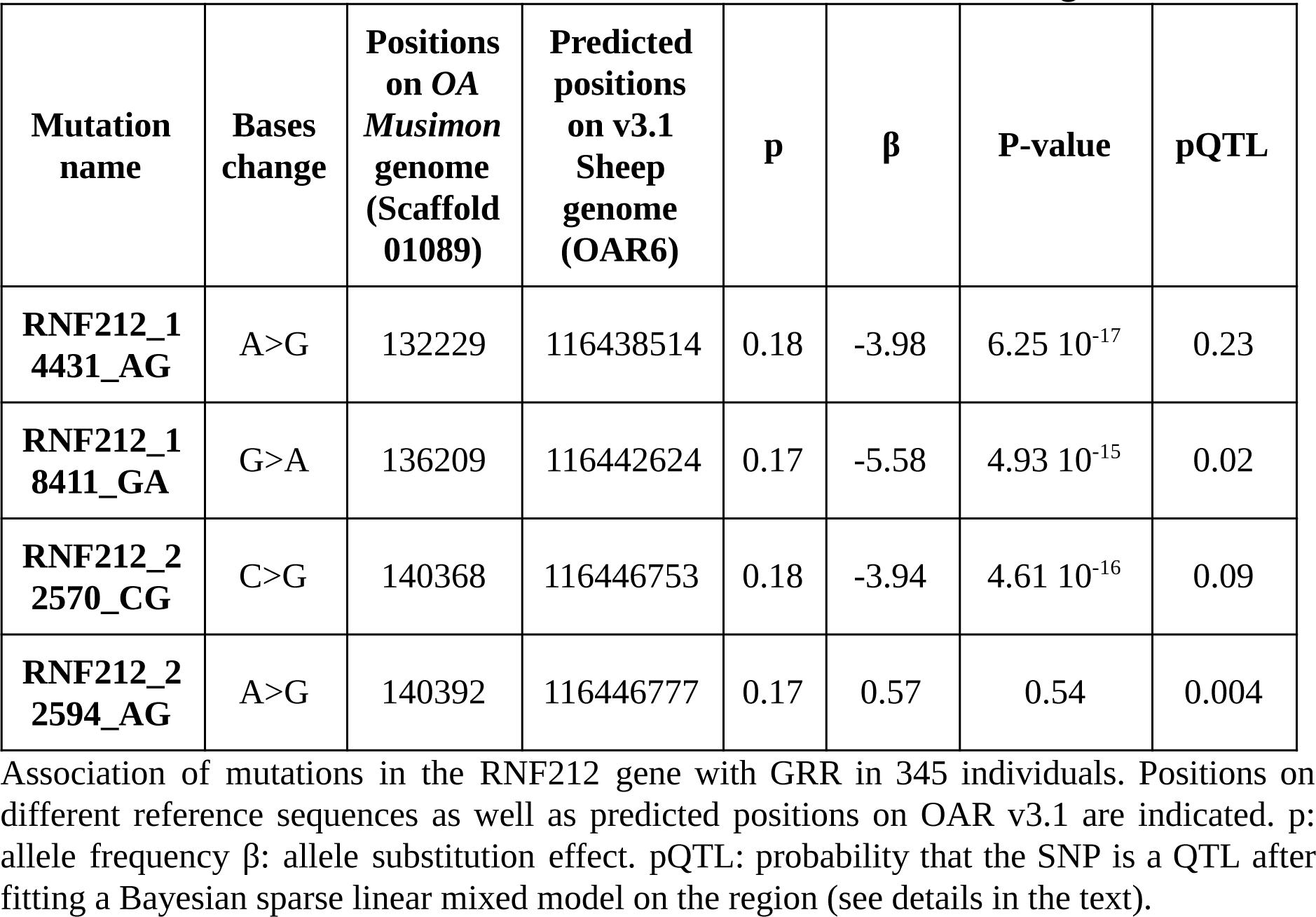
**Association of GRR with mutations in the *RNF212* gene.**

| **Mutation name** | **Bases change** | **Positions on OA *Musimon* genome (Scaffold 01089)** | **Predicted positions on v3.1 Sheep genome (OAR6)** | **p** | **β** | **P-value** | **pQTL** |
| --- | --- | --- | --- | --- | --- | --- | --- |
| **RNF212_1 4431_AG** | A>G | 132229 | 116438514 | 0.18 | -3.98 | 6.25 10<sup>-17</sup> | 0.23 |
| **RNF212_1 8411_GA** | G>A | 136209 | 116442624 | 0.17 | -5.58 | 4.93 10<sup>-15</sup> | 0.02 |
| **RNF212_2 2570_CG** | C>G | 140368 | 116446753 | 0.18 | -3.94 | 4.61 10<sup>-16</sup> | 0.09 |
| **RNF212_2 2594_AG** | A>G | 140392 | 116446777 | 0.17 | 0.57 | 0.54 | 0.004 |

Association of mutations in the *RNF212* gene with GRR in 345 individuals. Positions on different reference sequences as well as predicted positions on OAR v3.1 are indicated. p: allele frequency β: allele substitution effect. pQTL: probability that the SNP is a QTL after fitting a Bayesian sparse linear mixed model on the region (see details in the text).

## The genetic determinism of recombination differ between Soay and Lacaune males

GWAS in the Soay identified two major QTLs for GRR, with apparent sex-specific effects. These two QTLs were located in the same genomic regions as our QTLs on chromosome 6 and chromosome 7. The chromosome 6 QTL was only found significant in Soay females, while we detect a very strong signal in Lacaune males. Although the QTL was located in the same genomic region, the most significant SNPs were different in the two GWAS (Figure 6). Two possible explanations could be offered for these results: either the two populations have the same QTL segregating and the different GWAS hits correspond to different LD patterns between SNPs and QTLs in the two populations, or the two populations have different causal mutations in the same region. Denser genotyping data, for example by genotyping the RNF212 mutations identified in this work in the Soay population, would be needed to have a clear answer. For the chromosome 7 QTL, the signal was only found using regional heritability mapping (Nagamine et al. 2012) in the Soay, and after genotype imputation in our study, which makes it even more difficult to discriminate between a shared causal mutation or different causal mutations at the same location in the two populations.

## Discussion

In this study, we studied the distribution of recombination along the sheep genome and its relationship to historical recombination rates. We showed that contemporary patterns of recombination are highly correlated to the presence of historical hotspots. We showed that the recombination patterns along the genome are conserved between distantly related sheep populations but that their genetic determinism of genome-wide recombination rates differ. In particular, we showed that polymorphisms within the *RNF212* gene are strongly associated to male recombination in Lacaune whereas this genomic region shows no association in Soay males. Hence, combining three datasets, two pedigree datasets in distantly related domestic sheep populations and a densely genotyped sample of unrelated animals, revealed that recombination rate and its genetic determinism can evolve at short time scales, as we discuss below.

## Fine-scale Recombination Maps

In this work, we were able to construct fine-scale genetic maps of the sheep autosomes by combining two independent inferences on recombination rate. Our study on meiotic recombination from a large pedigree dataset revealed that sheep recombination exhibits general patterns similar to other mammals (Shifman et al. 2006; Chowdhury et al. 2009; Tortereau et al. 2012). First, sheep recombination rates were elevated at the chromosome ends, both on acrocentric and metacentric chromosomes. In the latter, our analysis revealed a clear reduction in recombination near centromeres. Second, recombination rate depended on the chromosome physical size, consistent with an obligate crossover per meiosis irrespective of the chromosome size. These patterns were consistent with those established in a very different sheep population, the Soay (Susan E. Johnston et al. 2016), and indeed when re-analysing the Soay data with the same approach as used in this study, the results showed a striking similarity between recombination rates in the two populations. Hence, our results show that recombination patterns were conserved over many generations and despite the very different evolutionary histories of the two populations and clear differences in the genetic determinism of GRR in males of the two populations. This similarity allowed to combine the two datasets to create more precise male sheep recombination maps than any of the two studies taken independently.

Our historical recombination maps revealed patterns of recombination at the kilobase scale, with small highly recombining intervals interspaced by more wide, low recombining regions. This result was consistent with the presence of recombination hotspots in the highly recombinant intervals. A consequence was that, as observed in other species, the majority of recombination took place in a small portion of the genome: we estimated that 80% of recombination takes place in 40% of the genome. (Kaur and Rockman 2014) suggested to use a Gini coefficient as a measure of the heterogeneity in the distribution of recombination along the genome to facilitate inter-species comparisons. When calculated on the historical recombination data, the Lacaune sheep has a coefficient of 0.52, which is similar to what is observed in Drosophila but lower than that measured in humans or mice. However, the coefficient calculated here is likely an underestimate due to our limited resolution (a few kilobases on the HD SNP array) compared to the typical hotspot width (a few hundred base-pairs). Overall, we identified 50,000 hotspot intervals which was twice the estimated number of hotspots in humans (International HapMap Consortium et al. 2007). This difference can be explained by different non mutually exclusive reasons. First, it is possible that what we detect as crossover hotspots are due to genome assembly errors and we indeed found a significant albeit moderate effect (OR 1.4) of the presence of assembly gaps in an interval on its probability of being called a hotspot. Second, our method to call hotspots could be too liberal. Indeed, a more stringent threshold (FDR=0.1%) would lead to about 25,000 hotspots, which would be similar to what is found in humans. Third, selection has been shown to impact hotspots discoveries although not with the methods we used here (Chan, Jenkins, and Song 2012). Finally, there exists the possibility that historically sheep exhibits more recombination hotspots than humans. In any case, the strong association between meiotic recombination rate and density in historical hotspots showed that our historical recombination maps were generally accurate. We tried to find enrichment in sequence motifs in the detected hotspots or specify their position relative to TSS (data not shown), but with no success mainly due to (i) the relative large hotspots intervals (about 5Kb) compared to typical hotspot motifs and (ii) the quality of the sheep genome assembly which still contains many small gaps that make such analyses difficult. Ultimately these questions would need an improved genome assembly and better resolution of crossover hotspots which should be addressed in the future from LD-based studies on resequencing data.

We combined, using a formal statistical approach, meiotic- and LD-based recombination rate estimates. Using an approach conceptually similar to that of (O’Reilly, Birney, and Balding 2008) led us to assess the impact of selection events on the sheep genome, in particular suggesting the possibility of an effect of diversifying selection at olfactory receptors genes. Based on this comparison, the correlation between historical and meiotic recombination rates was found to be high (r 0.7), but less than could be expected from previous results in humans, where the correlation was 97% on 5 Mb (S. Myers et al. 2006). However, it was closer to that of worms, mice or Drosophila, 69%, 47% and 50% respectively (Rockman and Kruglyak 2009, Brunschwig *et al.* 2012, Chan *et al.* 2012). Again, more precise estimates of both meiotic and historical recombination rates could change this number but other causes can be put forward.

A first explanation could come from the fact that the model we used to estimate historical recombination rates is based on the assumption of a constant effective population size, both in the past and along the genome. To allow for varying population size along the genome, we estimated the model in 2Mb intervals but there is still the possibility that varying population size in the past affect our historical recombination rate estimates, as the method has been shown to be somewhat influenced by demography although the identification of crossover hotspots much less so (N. Li and Stephens 2003). Also, as already mentioned above, selection has been shown to have substantial impact on estimation of recombination rates with other approaches (Chan, Jenkins, and Song 2012) although it has not been evaluated for the Li and Stephens model to our knowledge.

Second, our meiotic recombination maps are based on male meioses only while historical recombination rates are averaged over both male and female meioses. The fact that male and female recombination differ substantially, in particular in sheep (Susan E. Johnston et al. 2016), could also explain this relatively lower correlation.

Third, it is also possible that selective pressure due to domestication and later artificial breeding had the impact of modifying extensively LD patterns on the sheep genome, degrading the correlation between the two approaches. Indeed, the historical recombination estimates summarize ancestral recombinations that took place in the past and it is possible that recombination hotspots that were present in an ancestral sheep population are not longer active in today’s Lacaune individuals. This could arise, for example, if domestication led to a reduction in the diversity of hotspots defining genes, such as *PRDM9*, and hence a reduction in the number of motifs underlying hotspots which would in turn change the distribution of recombination on the genome. This has been shown for example in Humans were patterns of recombination differ between populations due to their different diversity at *PRDM9* (Berg et al. 2010, 2011; Baudat et al. 2010). Eventually, such a phenomenon would degrade the correlation between present day recombination (measured by the meiotic recombination rates) and past recombination (measured by historical recombination rates). Further studies on the determinism of hotspots in the sheep, its related genetic factors and their diversity would be needed to elucidate this question.

Despite these different effects, the substantial correlation between meiotic and historical recombination rates motivates the creation of scaled recombination maps that can be useful for interpreting statistical analysis of genomic data. As an illustration of the importance of fine-scale recombination maps for genetic studies, we found an interesting example in a recent study on adaptation of sheep and goat (Kim et al. 2016). In this study, a common signal of selection was found using the iHS statistic (Voight et al. 2006) in these two species (Figure 5 in (Kim et al. 2016)). This signature matches precisely the low recombining regions we identified on chromosome 10. However, the iHS statistic has been shown to be strongly influenced by variation in recombination rates, and in particular to tend to detect low recombining regions as selection signatures (Ferrer-Admetlla et al. 2014; O’Reilly, Birney, and Balding 2008). Precise genetic maps such as the one we provide in this work could thus help in annotating and interpreting such selection signals.

**Figure 5:**
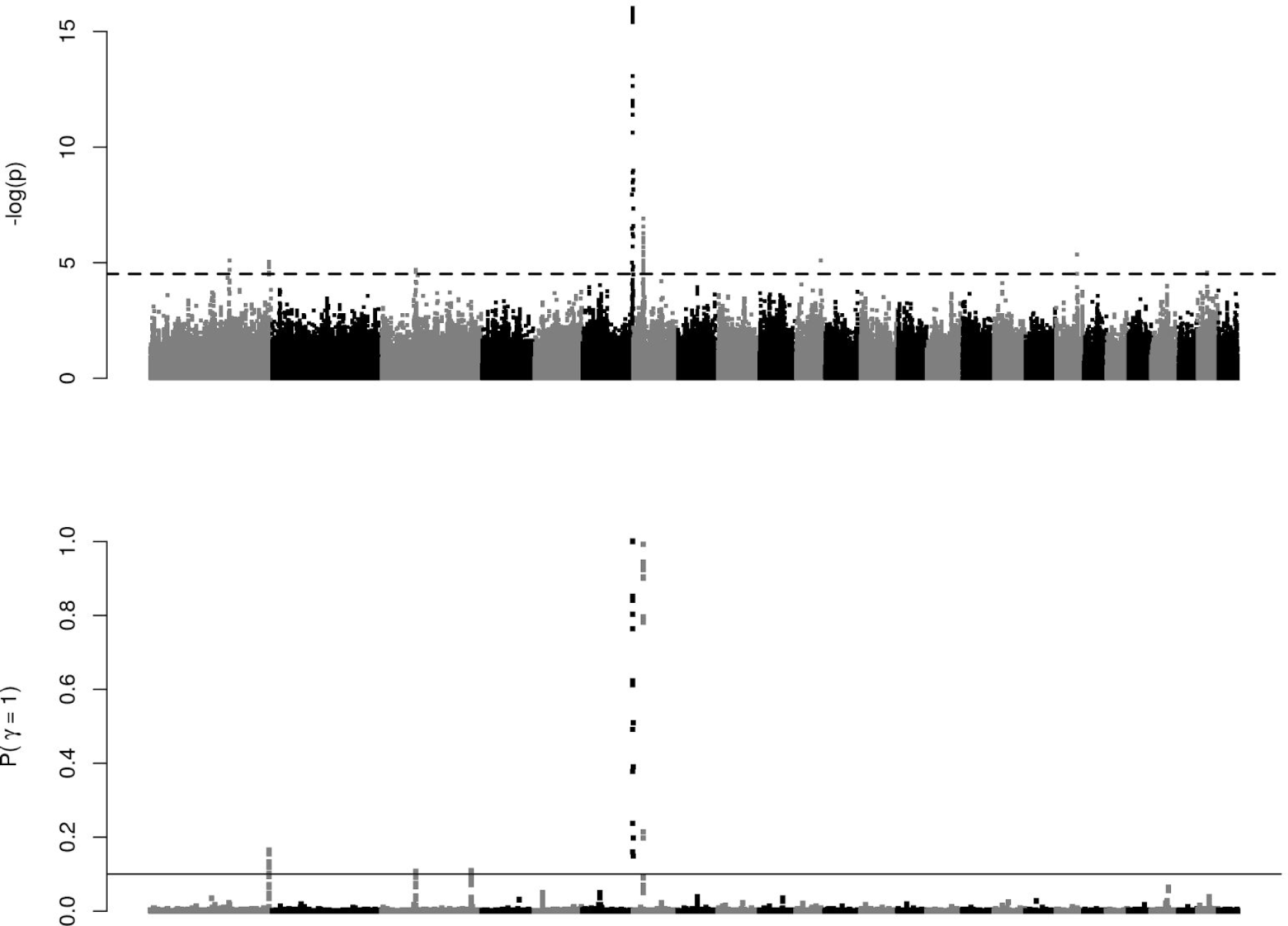
Genome-wide association study identifies three main QTLs for GRR. Top: −log10 (p-value) for single SNP tests for association. The genome-wide significance level (FDR=5%) is represented by the horizontal dotted line. Bottom: posterior probability that a region of 20 SNPs harbors a QTL, using a Bayesian multi-QTL model.

**Figure 6:**
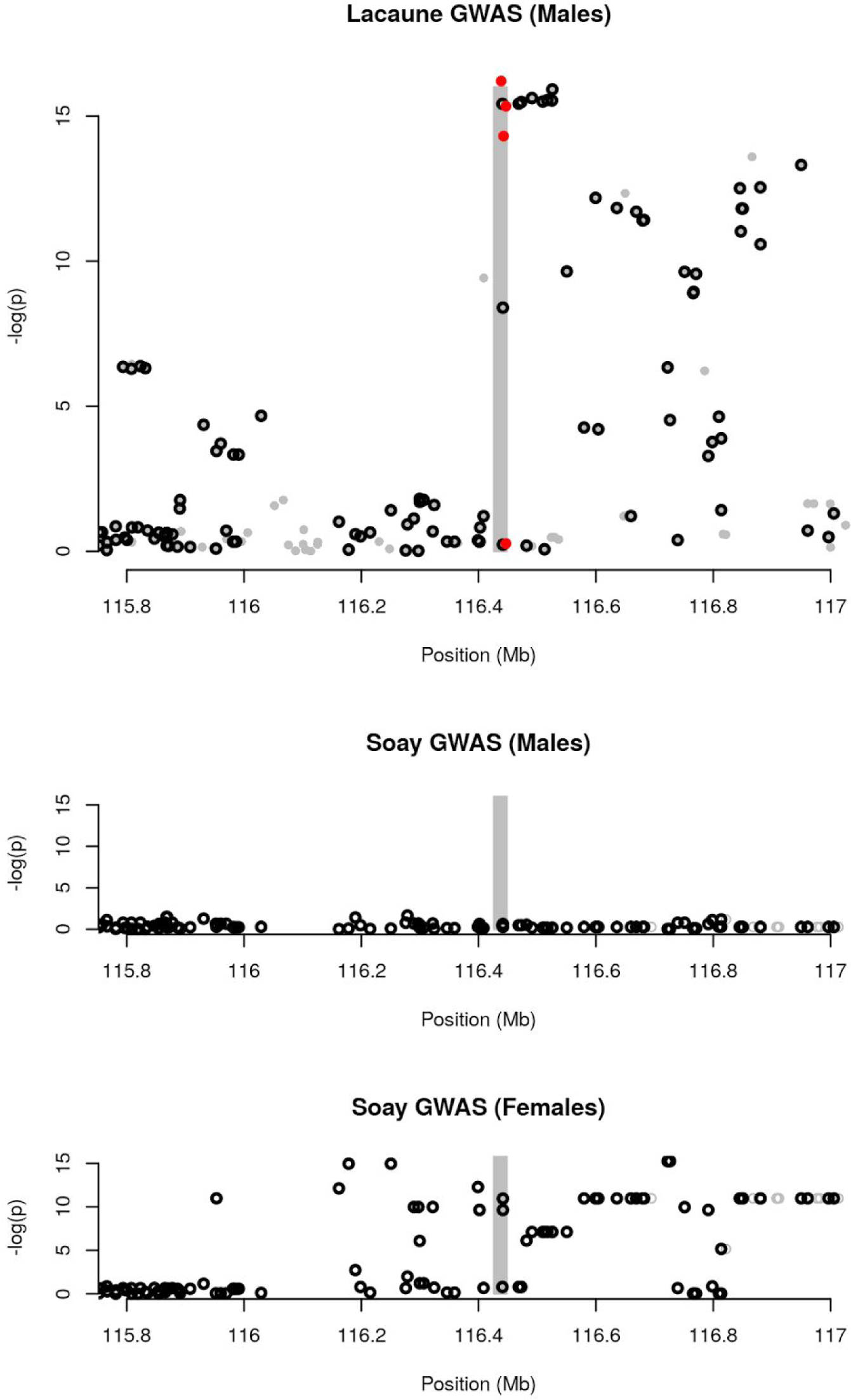
Comparison of GWAS results for the OAR 6 QTL in Lacaune Males (Top), Soay Males (Middle) and Soay Females (Bottom) The shaded area highlights the predicted position of the RNF212 gene. Circle dots are markers tested in both populations. Red dots are the new mutations within the RNF212 gene discovered in this study and genotyped in the Lacaune population. GWAS results in the Soay are from Johnston et al. (2016).

## Determinism of Recombination Rate in sheep populations

As mentioned in the introduction two phenotypes have been studies with respect to the recombination process, but only one was studied here, Genome-wide recombination rate (GRR). We found that our data was not sufficient to study the Individual Hotspot Usage, which requires either a larger number of meioses per individual (Ma et al. 2015; Kadri et al. 2016a; Sandor et al. 2012a) or denser genotyping in families (Coop et al. 2008).

Our approach to study the genetic determinism of GRR in the Lacaune population was first to estimate its heritability, using a classical analysis in a large pedigree. This analysis also allowed to extract additive genetic values (EBVs) for the trait in 345 male parents, which we used for a GWAS in a second step. The EBVs are by definition, only determined by genetic factors, as environmental effects on GRR are averaged out. Indeed, we found that the proportion of variance in EBVs explained by genetic factors in the GWAS was essentially one. A consequence was that, although this sample size could be deemed low in current standards, the power of our GWAS was greatly increased by the high precision on the phenotype. We estimated the heritability of GRR at 0.23, which was similar to estimates from studies on the same phenotype in ruminants (*e.g.* 0.22 in cattle (Sandor et al. 2012a) or 0.12 in male Soay sheep (Susan E. Johnston et al. 2016), but see below for a discussion on the comparison with Soay sheep). We had little information on the environmental factors that could influence recombination rate, but did find a suggestive effect of the month of insemination on GRR, especially we found increased GRR at the month of May. Confirmation and biological interpretation of this result would need dedicated studies, but it was consistent with the fact that fresh (*i.e.* not frozen) semen is used for insemination in sheep and that the reproduction of this species is seasonal (Rosa and Bryant 2003).

The genetic determinism of GRR discovered in our study closely resembles what has been found in previous studies, especially in mammals. Two major loci and two suggestive ones affected recombination rate in Lacaune. The two main QTLs are common to cattle and Soay sheep. The underlying genes and mutations for these two QTLs are not yet resolved but the fact that the two regions harbour interacting genes (*RNF212* and *HEI10* (Qiao et al. 2014; Rao et al. 2016)) involved in the maturation of crossovers, make these two genes likely functional candidates. Indeed, these two genes were identified as potential candidates underlying QTLs for GRR in mice (R. J. Wang and Payseur 2017). The third gene identified here, *KCNJ15*, is a novel candidate, and its role and mechanism of action in the repair of DSBs needs to be confirmed and elucidated. Interestingly, these three genes are linked to the reparation of DSBs and crossover maturation processes. Finally, the fourth candidate *FSHR* has well documented effects on gametogenesis but has not been linked to recombination previously.

In our study, sixty percent of the additive genetic variance in GRR remained unexplained by large effect QTLs and were due to polygenic effects. This could be interpreted in the light of recent evidence that has shown that other mechanisms, involved in chromosome conformation during meiosis, explain a substantial part of the variation in recombination rate between mouse strains (Baier et al. 2014) and bovids (Ruiz-Herrera et al. 2017). Furthermore the variations at the major mammal recombination loci (*RNF212*, *CPLX1*, *REC8* or the Human inversion *17q21.31*) explain only 3 to 11% (Ritz, Noor, and Singh 2017) of the phenotypic variance among individuals. Elucidating the genetic determinism of these different processes would thus require much larger sample sizes or different experimental approaches (Baier et al. 2014; Ruiz-Herrera et al. 2017).

The combination of datasets from the Lacaune population and one from the recent study of recombination in Soay sheep (Susan E. Johnston et al. 2016) allowed to study the evolution of recombination at relatively short time scales. One of the most striking difference between our two studies is that the two QTLs that were detected in common had no effect in Soay males, whereas they had strong effects in Lacaune males. However, the two populations had very similar polygenic heritability: accounting for the fact that the Lacaune QTLs explain about 40% of the additive genetic variance, we could estimate the polygenic additive genetic variance in Lacaune males at 0.16, very similar to the 0.12 found in Soay males. Combined with our results that the two populations exhibit very similar male recombination maps, both in terms of intensity and genome distribution, the combination of the two studies shows that recombination patterns are conserved between populations under distinct genetic determinism, highlighting the robustness of mechanisms that drive them. Further work is needed to get a more detailed picture of the genetic control of recombination in sheep and will likely require combining multiple inferences from genetics, cytogenetics, molecular biology and bioinformatics analyses.

## Acknowledgments

Institut de l’Elevage (JM. Astruc) and breed organizations (Ovitest and Confédération de Roquefort) provided the SNP genotypes and pedigree information. This work was partially funded by the BoDeliRe grant of the INRA Selgen Metaprogram and by Région Midi-Pyrénées. We are thankful to Tom Druet, Laurent Duret, Alain Pinton and Pierre Sourdille for their helpful comments on the manuscript.

---

**Figure S1.**
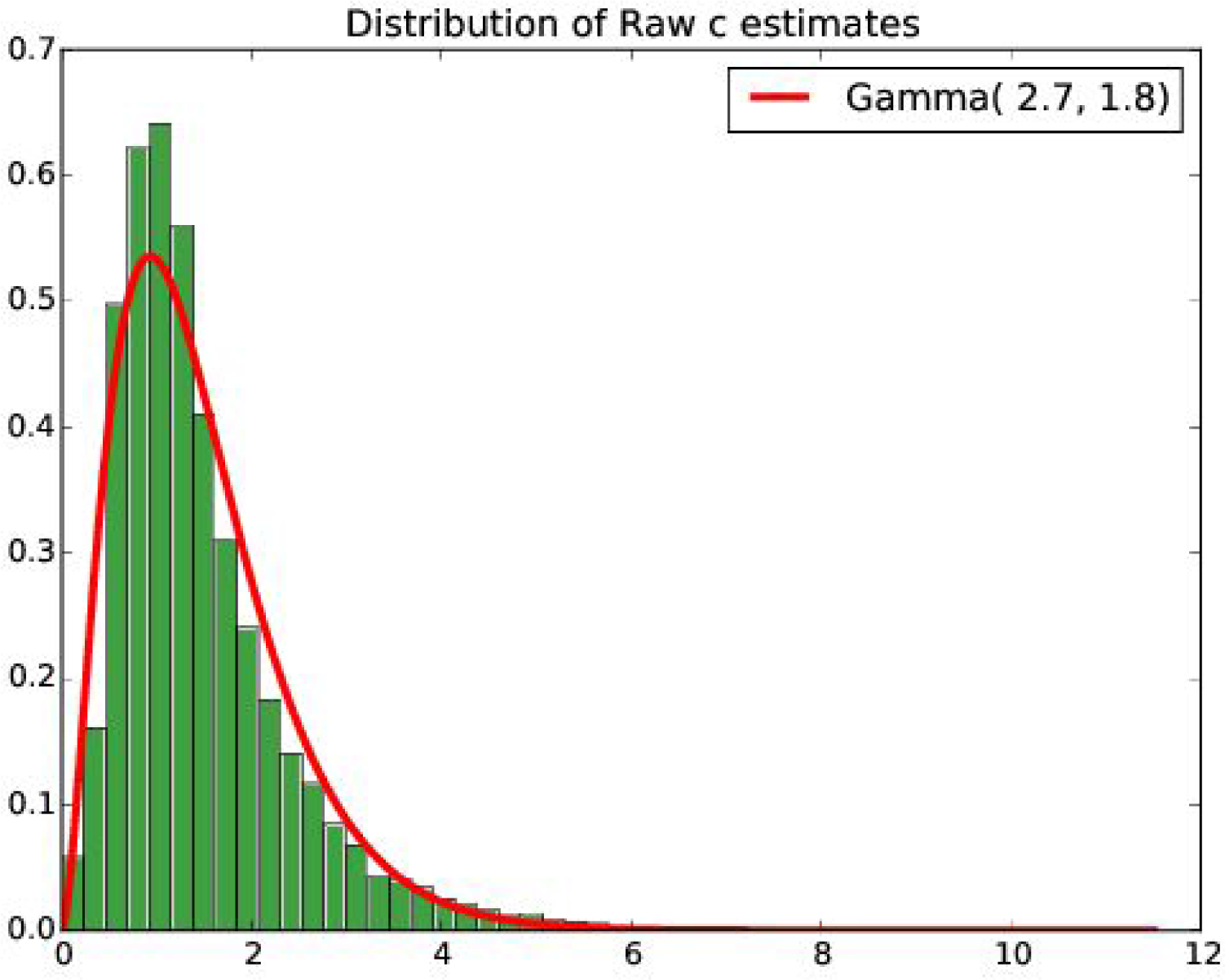
Genomewide distribution of recombination rate using the approach of Sandor et al. (2012). These estimates were calculated from observed crossover frequencies. Fitting a gamma distribution on the observed estimates (red line) provided a prior distribution for the subsequent Bayesian inference on recombination rates (parameters given in the box). See methods for details.

**Figure S2.**
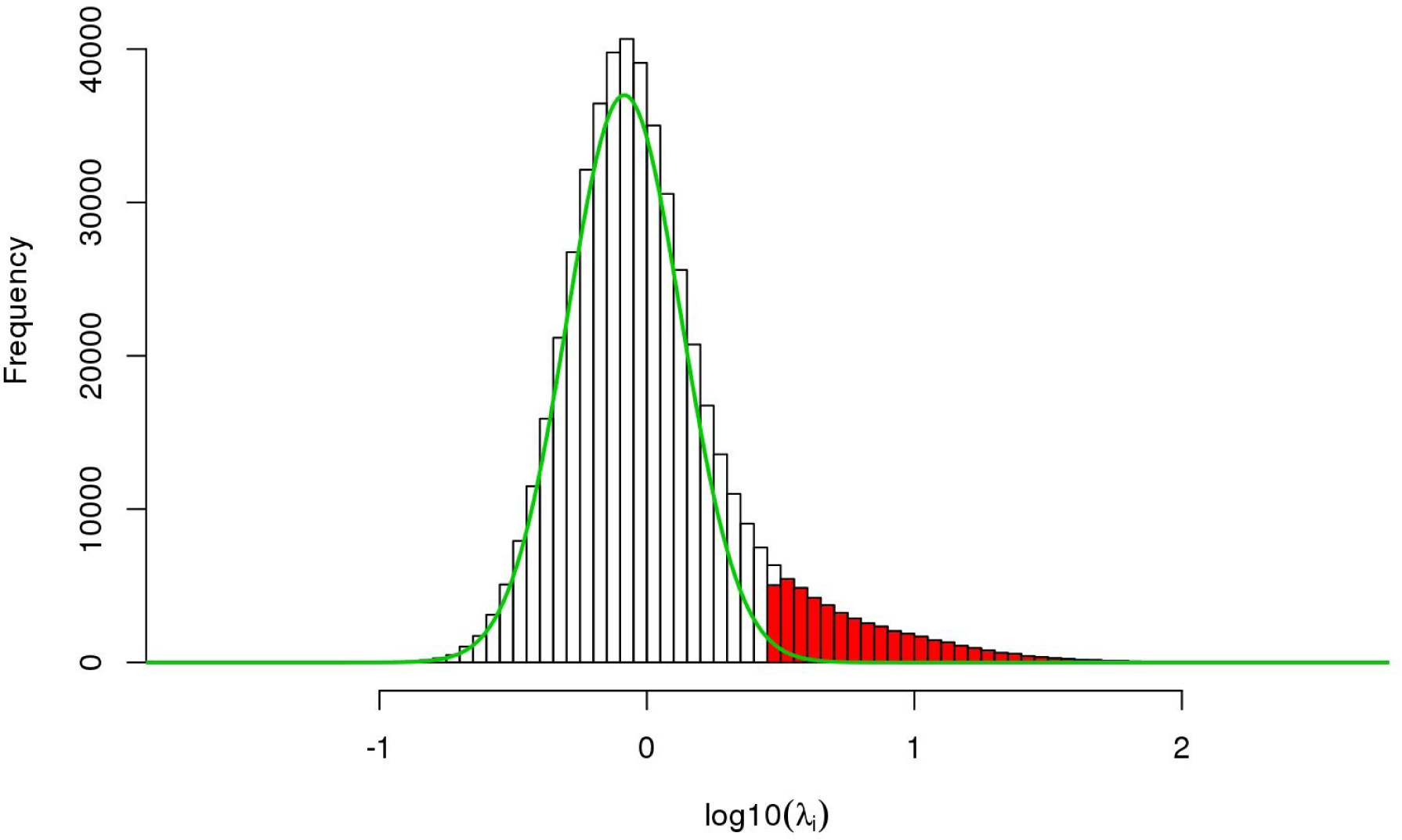
Distribution of recombination intensities among intervals of the HD SNP array. The green curve represents the distribution under the null hypothesis that there is no hotspots (log10(λ_i_) = 0). It is estimated by fitting a mixture of Gaussian distribution to the observed distribution and extracting the relevant component. Intervals where recombination intensity was particularly high (FDR = 5%) were considered as harbouring recombination hotspots and are shown in red.

**Figure S3.**
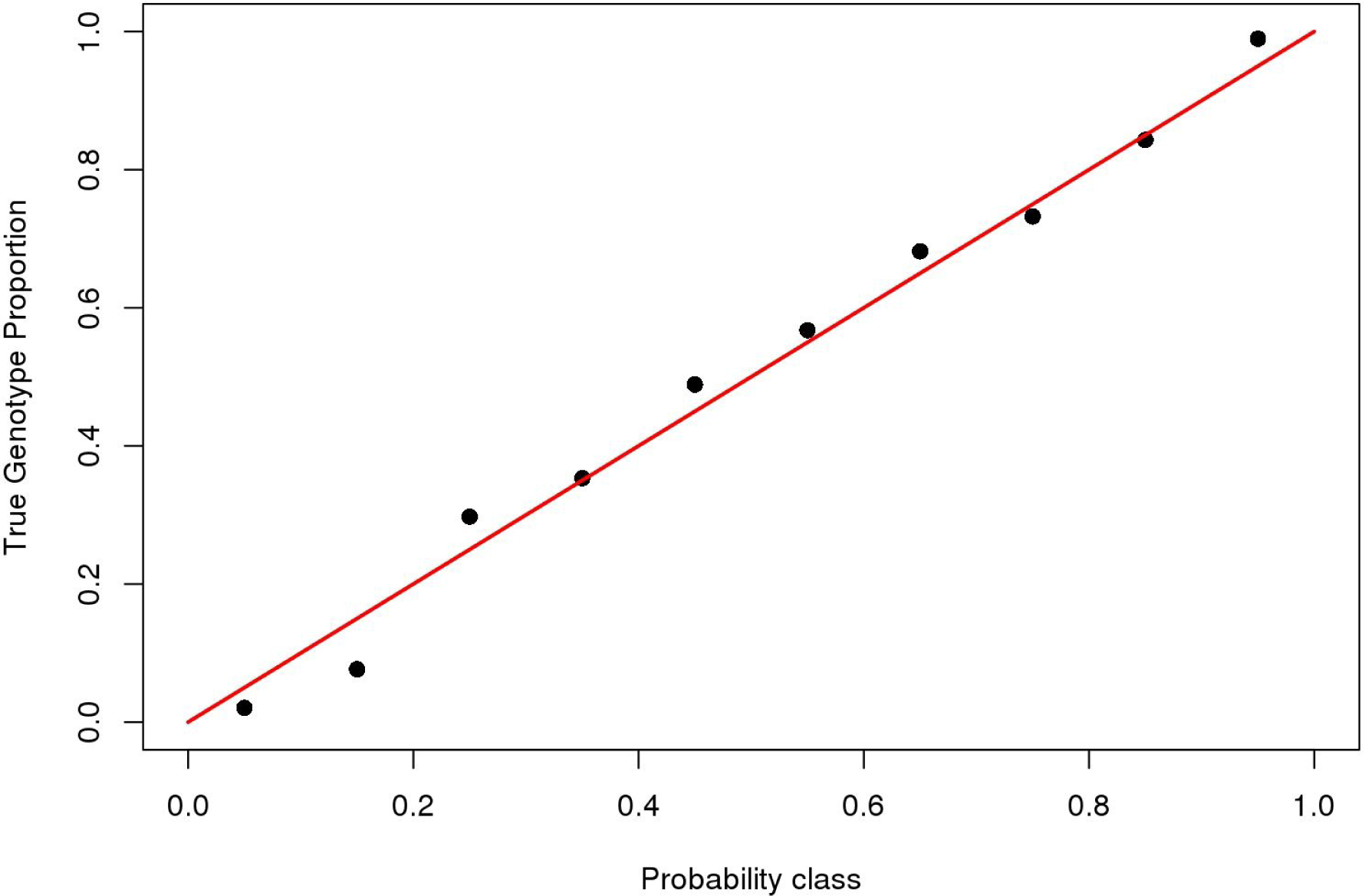
Validation of imputed genotypes for the GWAS. The figure shows the proportion of correct genotype calls as a function of their posterior probability calculated with BIMBAM.

**Figure S4.**
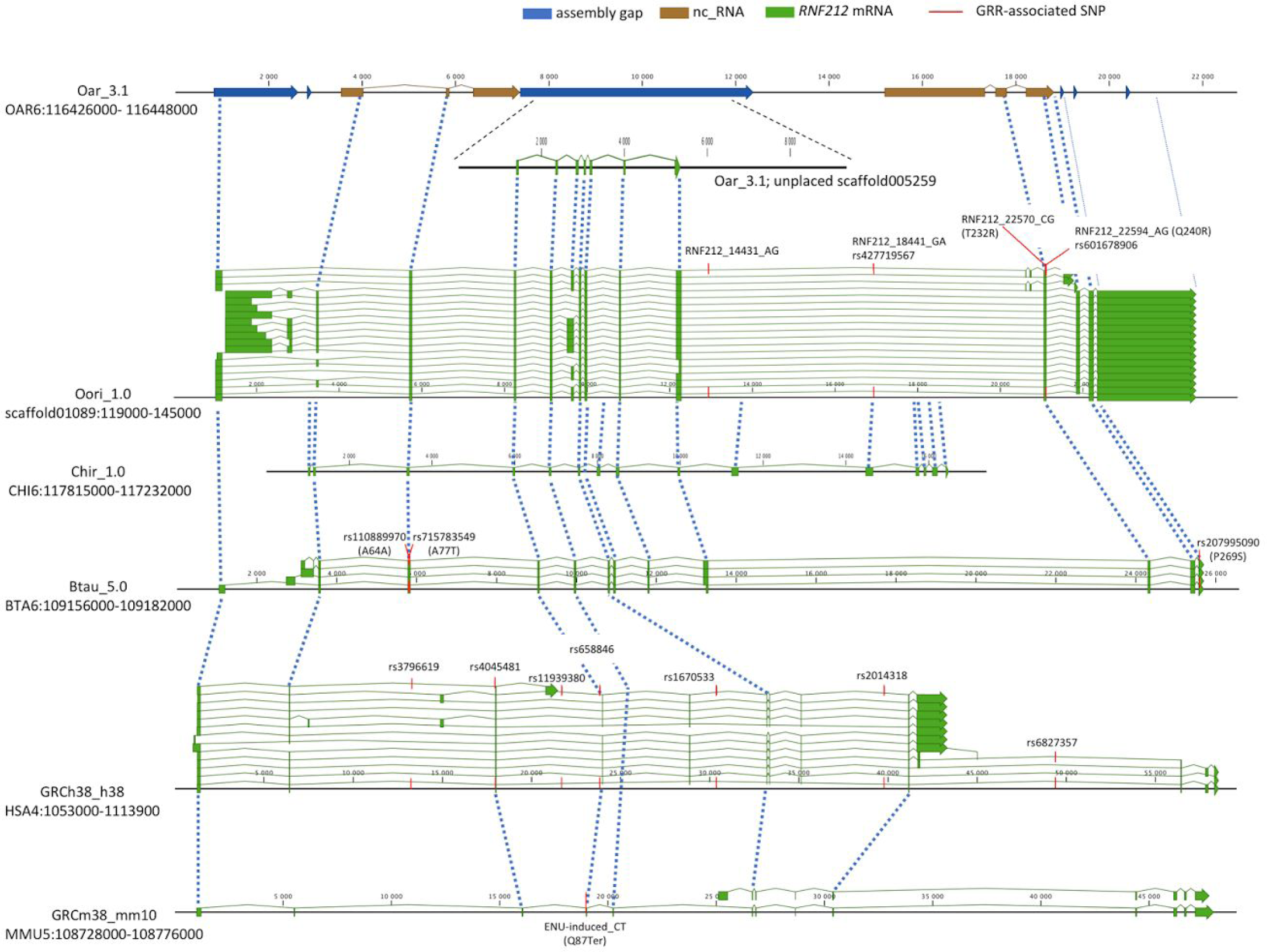
*RNF212* gene structure in various species. *RNF212* gene is not annotated on the ovine reference genome Oar_v3.1, but can be located at the telomeric end of OAR6 (116,4Mb) by homologies (dashed lines) with the *RNF212* gene from Ovis aries musinon (Oor1_1.0). Some anotated predicted non-coding RNA sequence (nc_RNA in brown) were part of the *RNF212* sequence. The ovine *RNF212* gene is also partly present in the unplaced scaffold005259, that can be virtually located in the largest assembly gap (in blue). In Ovis orientalis musimon, the *RNF212* gene exibited 14 putative exons with alternative splicing (mRNA models in green). Homology analysis (dashed lines) with annotated *RNF212* gene in other ruminant species (bovine on BTA6 and caprine on CHI6) indicated a good gene structure conservation between ovine and bovine *RNF212*, but only a partial conservation with goat *RNF212*, where the six last predicted exons match with intronic region in ovine. When compared to human *RNF212* on HSA4 and mouse *RNF212* on MMU5 chromosomes, only four to five exons are conserved with ruminants indicated a non-conserved gene structure. Red lines located SNP associated with global recombination rate (GRR) in the present study, and those previously shown in bovine (Sandor et al. 2012; Kadri et al. 2016), in human (Kong et al. 2008; Chowdhury et al. 2009; Fledel-Alon et al. 2011; Kong et al. 2014) and in mouse (Fujiwara et al. 2014). Gene scales are in base pair and gene structures were constructed with CLC Main Workbench software v7.7.3 using the NCBI query module (Qiagen Aarhus).

**Figure S5.**
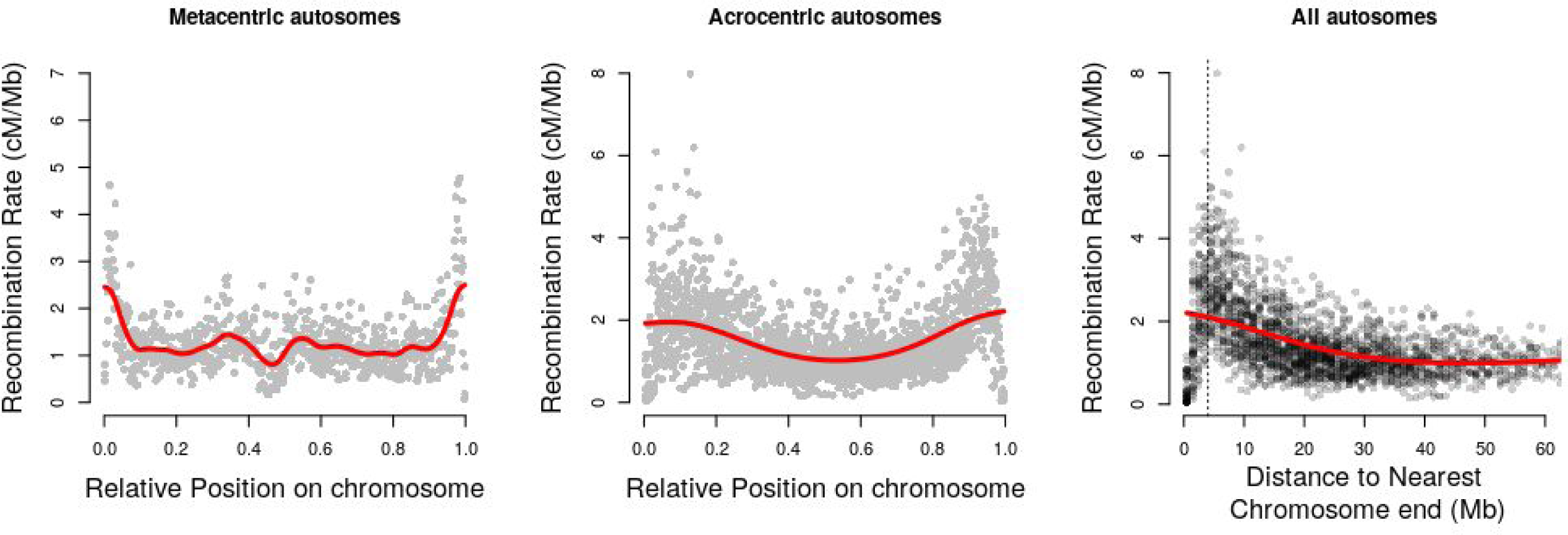
Patterns of recombination along Sheep autosomes. Left: recombination rate of one megabase windows along metacentric chromosomes (1,2,3). Center: recombination rate of one megabase windows along acrocentric chromosomes (4-26). Right: recombination rate of one megabase windows against distance to nearest chromosome end.

**Figure S6.**
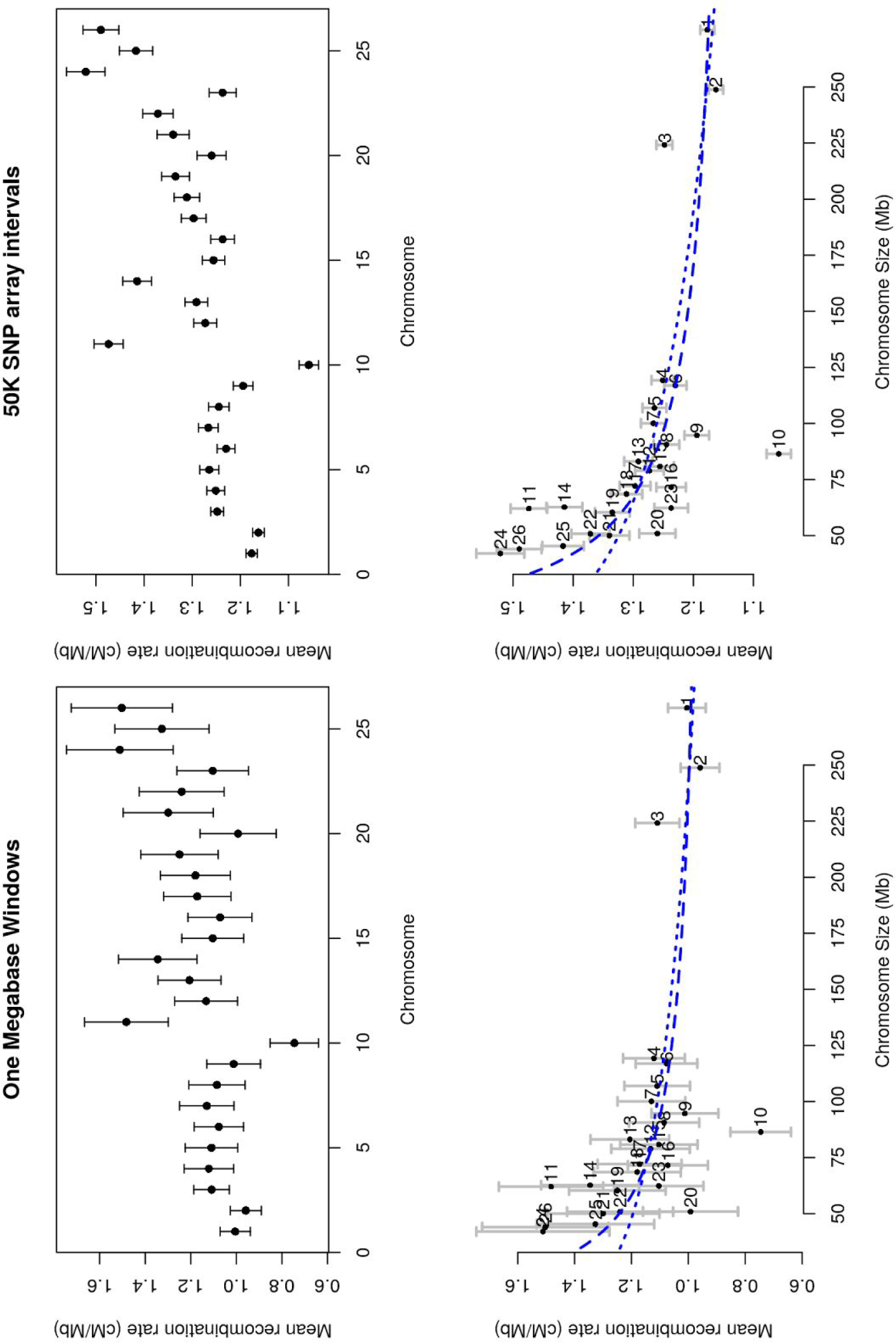
Recombination rates of Sheep autosomes. Left: from recombination rate estimates in windows of one megabase. Right: from recombination rate estimates in SNP array intervals. Top: for each chromosome. Bottom: as a function of chromosome physical size. Dotted line: c = f(log(size)), dashed line: c = f(1/size).

**Figure S7.**
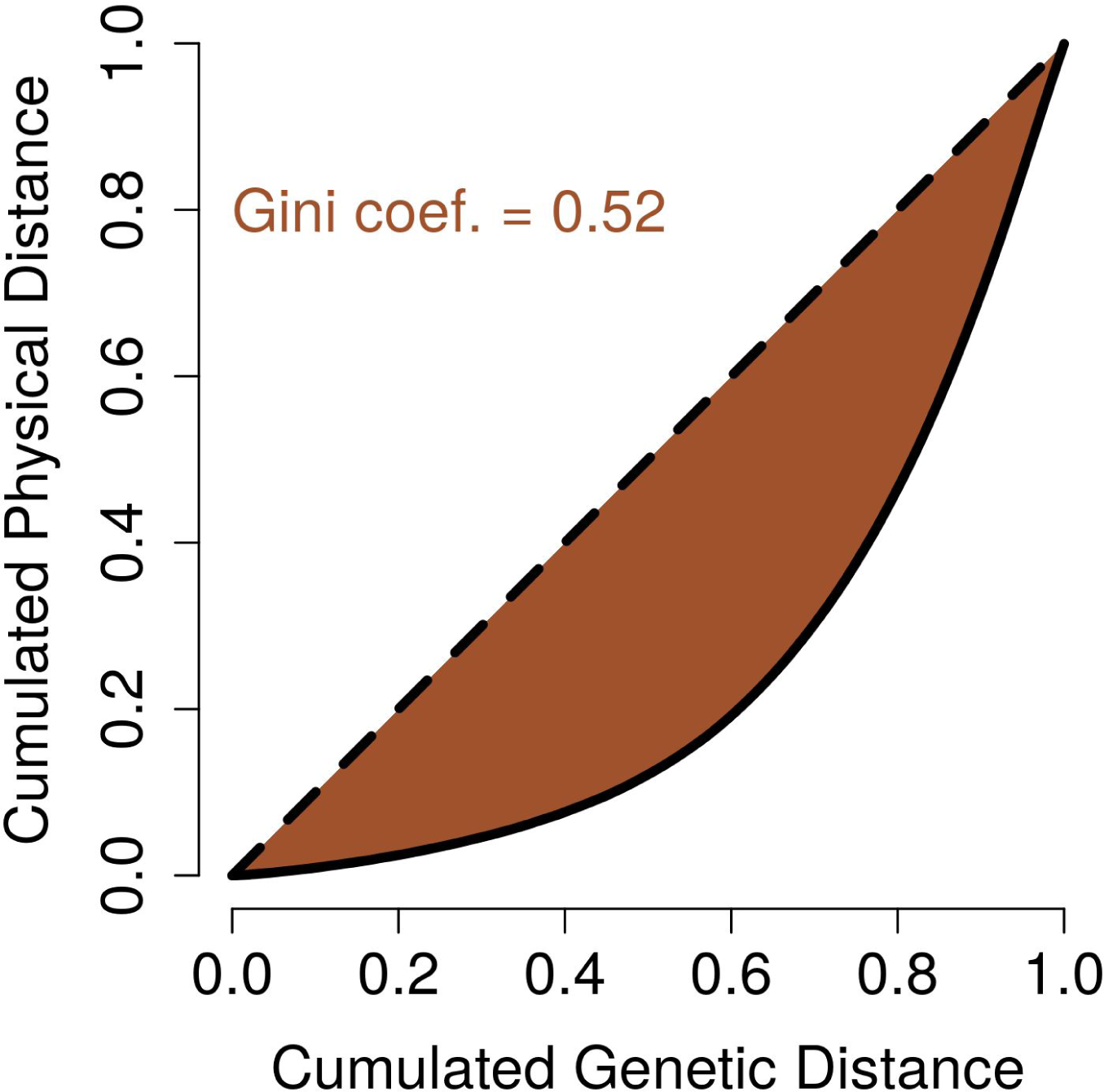
Distribution of recombination on the genome. The figure represents the proportion of the physical genome size affected by recombination, for increasing coverage of the genetic map. The Gini corresponding to the brown area on the figure is 0.52.

**Figure S8.**
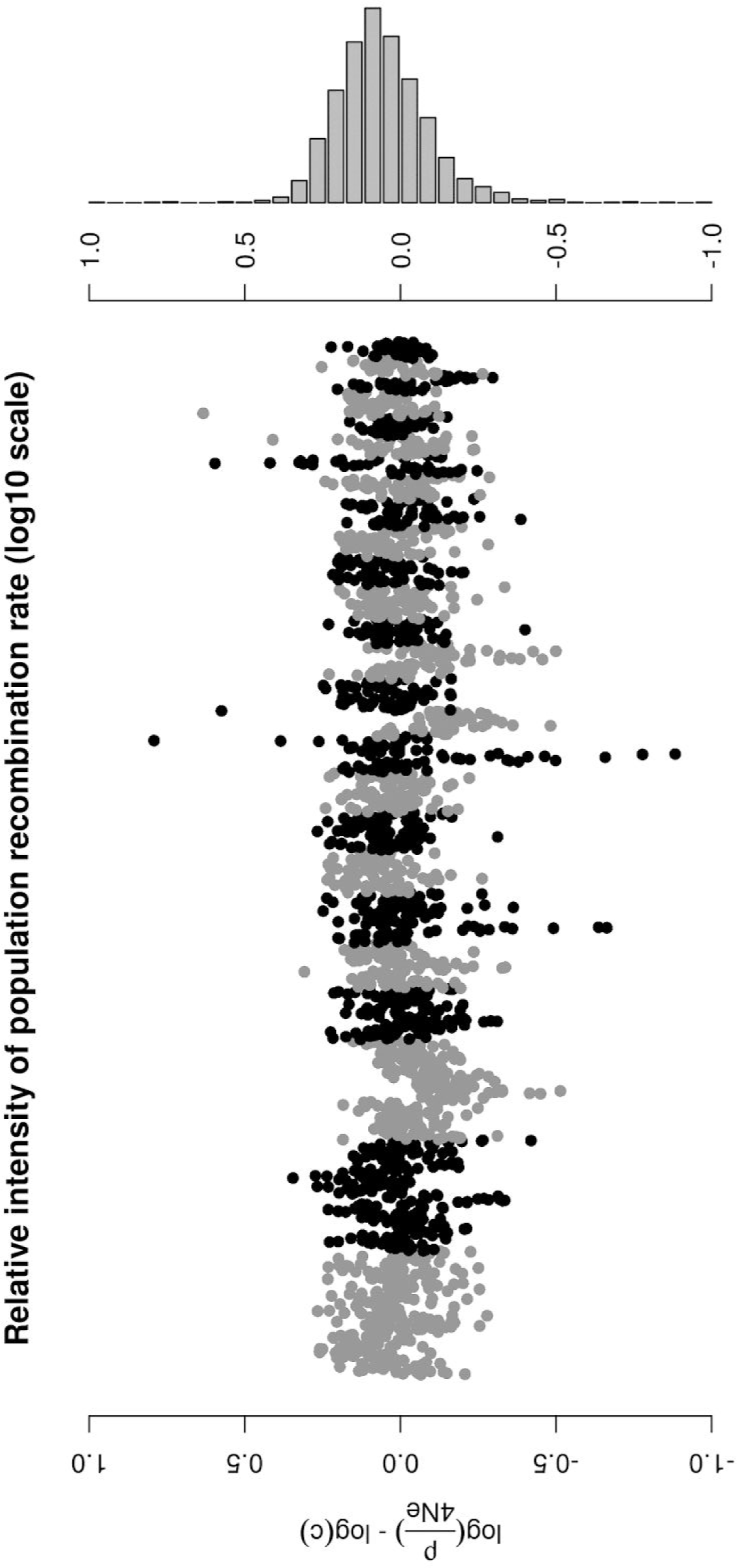
Relative intensity of population to meiotic recombination rates in windows of 1 Mb along the sheep genome.

**Figure S9.**
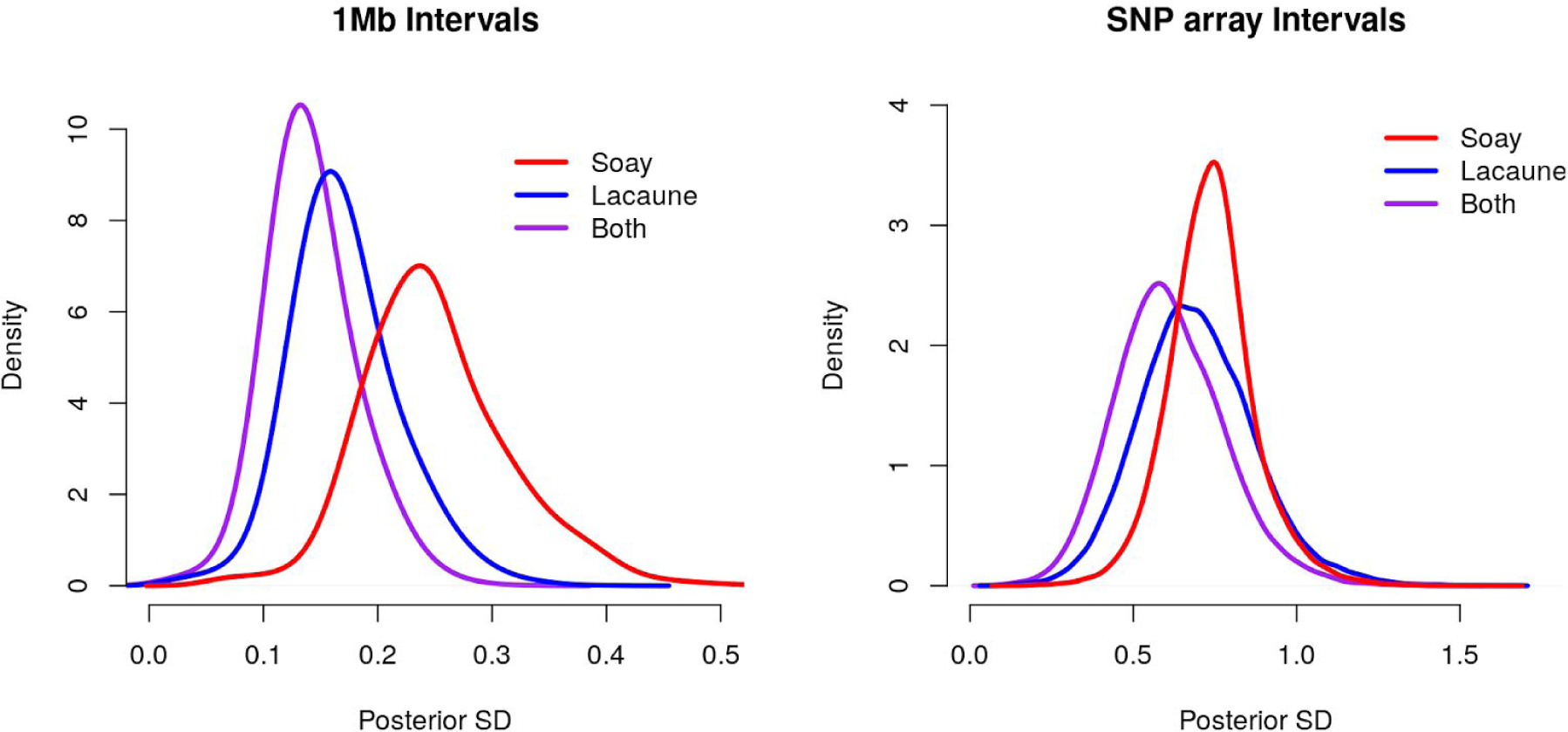
Posterior standard deviation of recombination rates on the medium density SNP array with different datasets. Soay: dataset from Johnston et al. (2016), only male meioses were used. Lacaune: dataset from this study. Both: combination of the two datasets.

**Figure S10.**
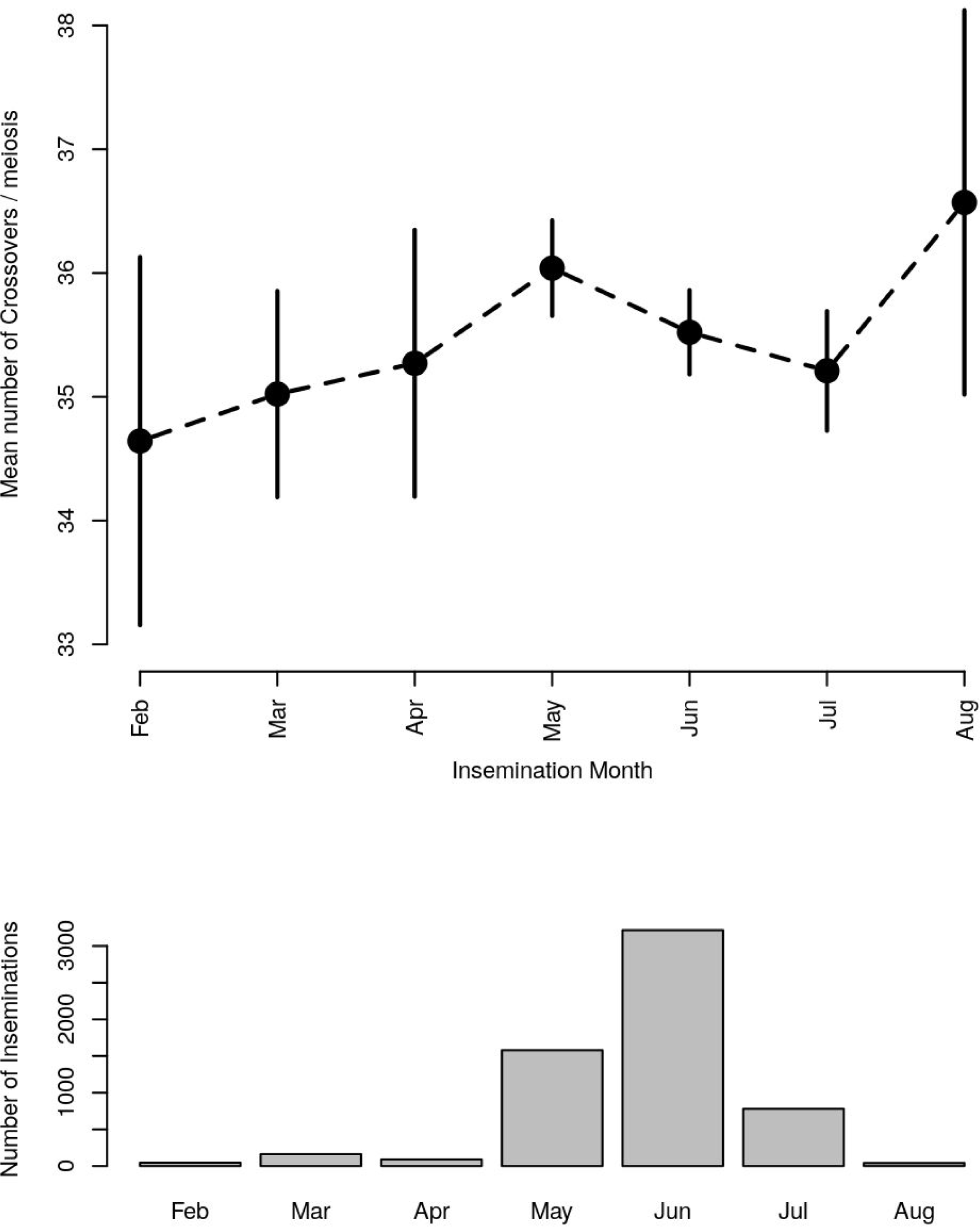
Effect of insemination month on the average number of crossovers per meiosis. Top: estimated mean GRR (dots) and 95% confidence intervals (vertical lines). Bottom: number of inseminations per month.

**Figure S11.**
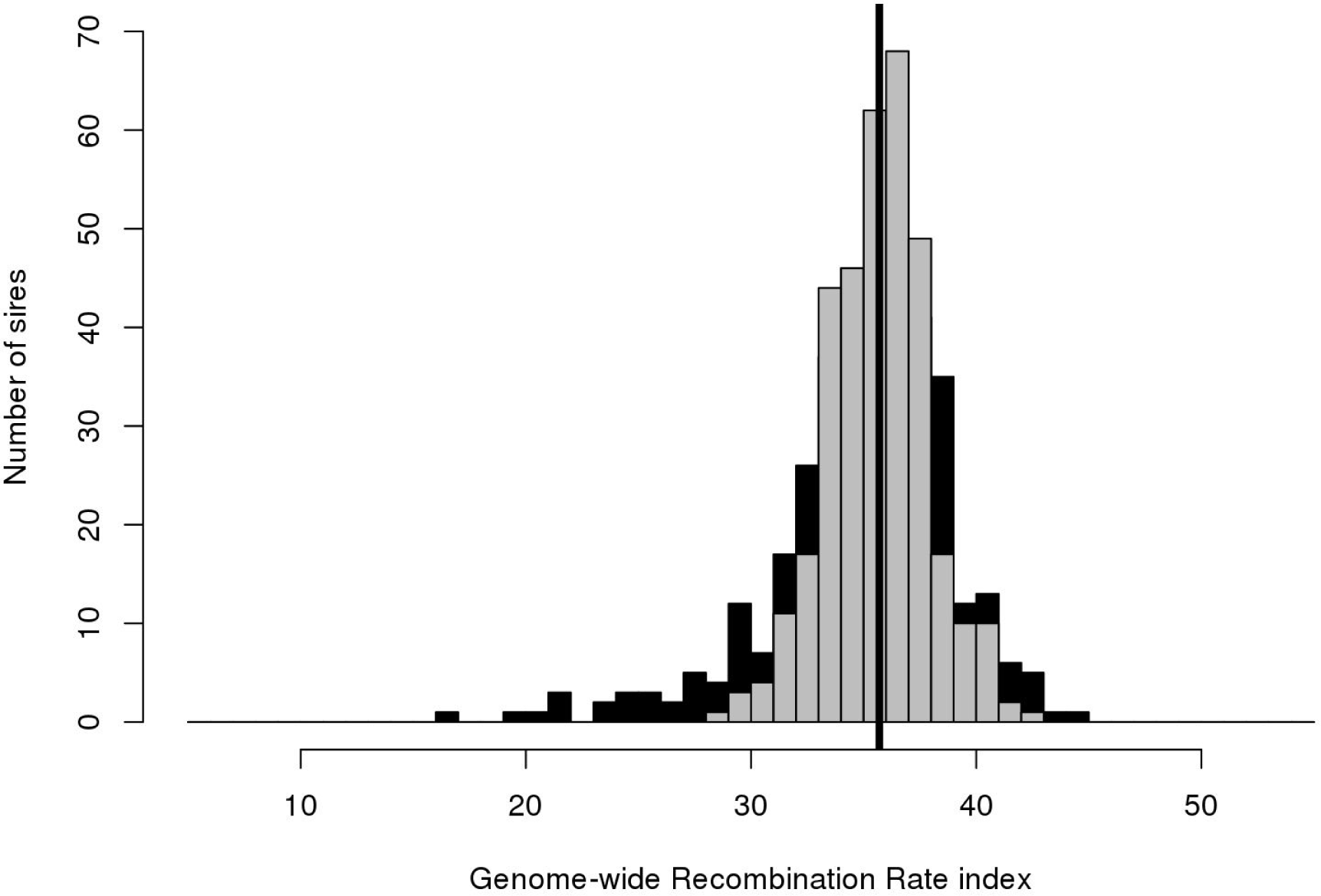
Individual variation in recombination rates among Lacaune Males. Additive genetic values on Genome-wide Recombination Rate genetic for all Lacaune sires of our dataset (in black) and for the 345 FID (in grey). The vertical black line is placed at the mean.

**Figure S12.**
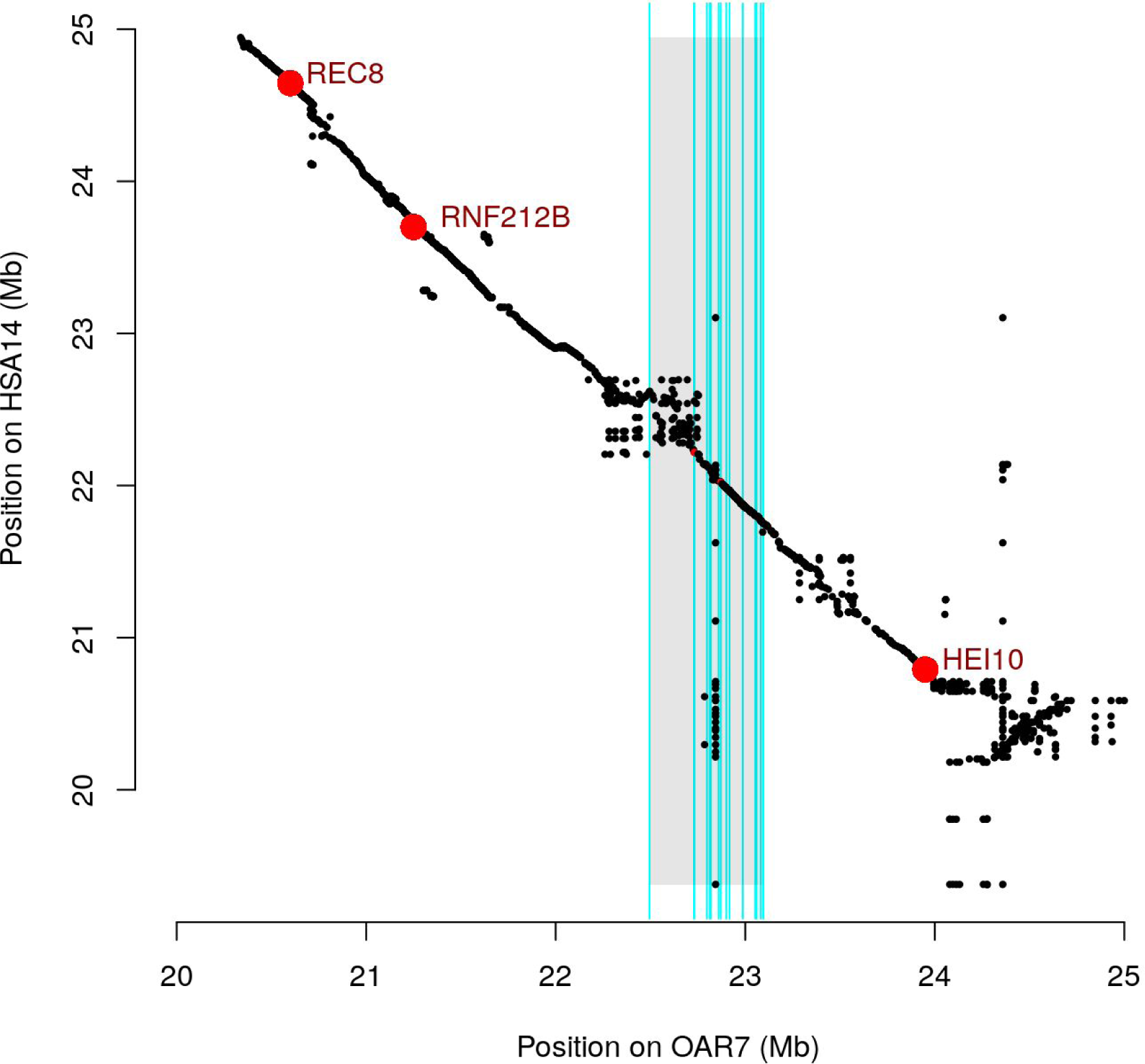
Local alignments of the Sheep and Human genome around the OAR7 QTL region. Dotplot of the alignements of sheep OAR7 on human HSA14. Vertical cyan bars are located at significant SNP positions. Three functional candidate genes surrounding the association signal (shaded) are indicated.

**Figure S13.**
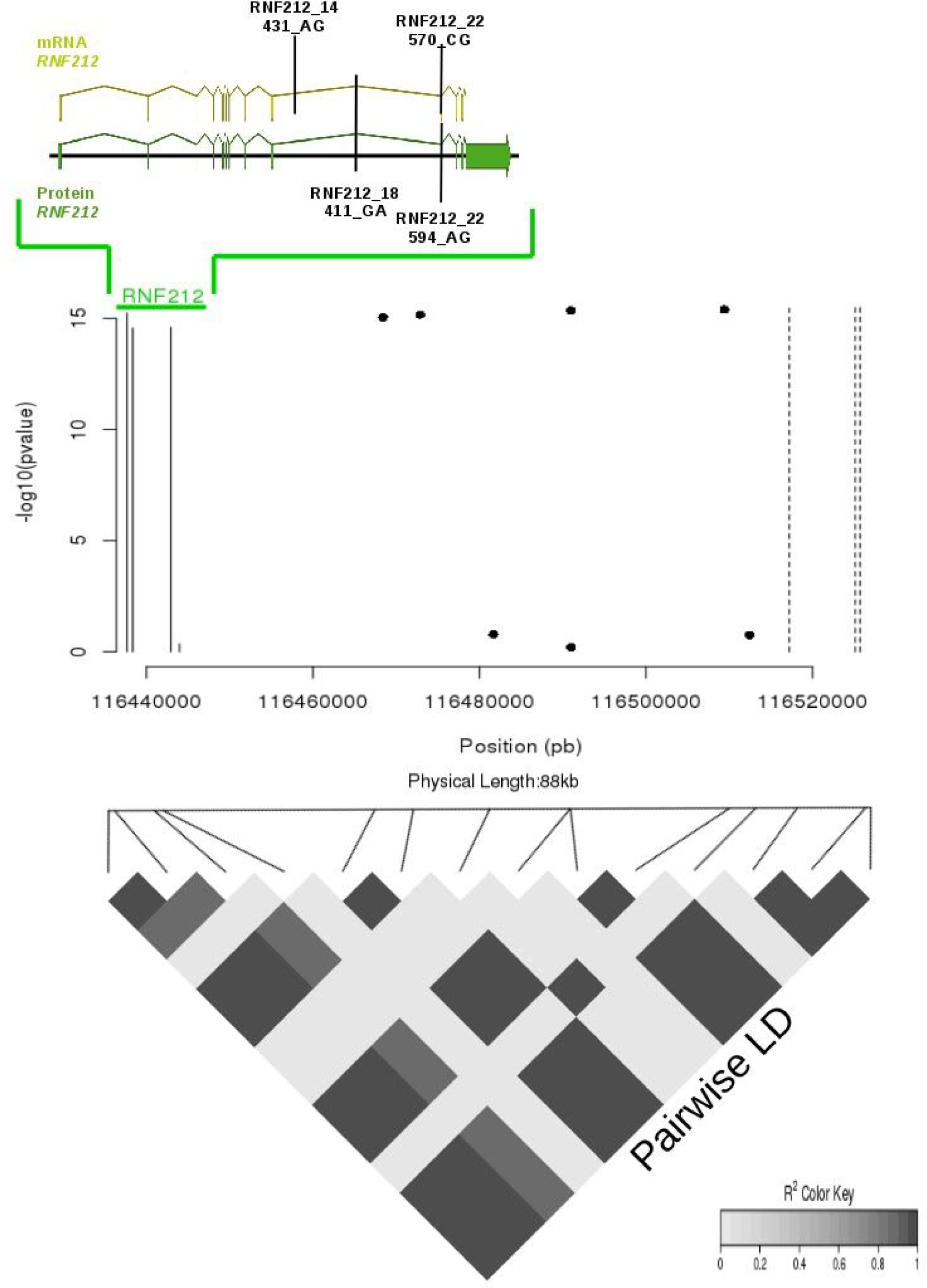
Linkage disequilibrium between *RNF212* polymorphisms and chromosome 6 QTL SNPs. The top figure represents the mRNA and the protein of *RNF212*. The four genotyped mutations are indicated: the two first are intronic and the two others are exonic. We replace the gene on a zoom on the chromosome 6 QTL (middle figure). The four left solid lines highlight the mutations, whereas the dashed lines represent the 3 most significant SNPs. Middle points show the intermediate SNPs between the mutations and the significant SNPs. Finally, the figure at the bottom indicates the pairwise LD between the mutations and all the SNPs presented on the middle figure. It highlights two haplotype blocks: one between the 3 most significant mutations and another between the 3 most significant SNPs.

## Literature Cited

Ahlawat, S., P. Sharma, R. Sharma, R. Arora, N. K. Verma, B. Brahma, P. Mishra, and S. De. 2016. “Evidence of Positive Selection and Concerted Evolution in the Rapidly Evolving PRDM9 Zinc Finger Domain in Goats and Sheep.” Animal Genetics 47 (6): 740–51.

Auton, Adam, Ying Rui Li, Jeffrey Kidd, Kyle Oliveira, Julie Nadel, J. Kim Holloway, Jessica J. Hayward, et al. 2013. “Genetic Recombination Is Targeted towards Gene Promoter Regions in Dogs.” PLoS Genetics 9 (12): e1003984.

Baier, Brian, Patricia Hunt, Karl W. Broman, and Terry Hassold. 2014. “Variation in Genome-Wide Levels of Meiotic Recombination Is Established at the Onset of Prophase in Mammalian Males.” PLoS Genetics 10 (1): e1004125.

Baloche, G., A. Legarra, G. Sallé, H. Larroque, J-M Astruc, C. Robert-Granié, and F. Barillet. 2014. “Assessment of Accuracy of Genomic Prediction for French Lacaune Dairy Sheep.” Journal of Dairy Science 97 (2): 1107–16.

Bates, Douglas, Martin Mächler, Ben Bolker, and Steve Walker. 2015. “Fitting Linear Mixed-Effects Models Using lme4.” Journal of Statistical Software 67 (1): 1–48.

Baudat, F., J. Buard, C. Grey, A. Fledel-Alon, C. Ober, M. Przeworski, G. Coop, and B. de Massy. 2010. “PRDM9 Is a Major Determinant of Meiotic Recombination Hotspots in Humans and Mice.” Science 327 (5967): 836–40.

Berg, Ingrid L., Rita Neumann, Kwan-Wood G. Lam, Shriparna Sarbajna, Linda Odenthal-Hesse, Celia A. May, and Alec J. Jeffreys. 2010. “PRDM9 Variation Strongly Influences Recombination Hot-Spot Activity and Meiotic Instability in Humans.” Nature Genetics 42 (10): 859–63.

Berg, Ingrid L., Rita Neumann, Shriparna Sarbajna, Linda Odenthal-Hesse, Nicola J. Butler, and Alec J. Jeffreys. 2011. “Variants of the Protein PRDM9 Differentially Regulate a Set of Human Meiotic Recombination Hotspots Highly Active in African Populations.” Proceedings of the National Academy of Sciences of the United States of America 108 (30): 12378–83.

Boitard, Simon, Willy Rodríguez, Flora Jay, Stefano Mona, and Frédéric Austerlitz. 2016. “Inferring Population Size History from Large Samples of Genome-Wide Molecular Data - An Approximate Bayesian Computation Approach.” PLoS Genetics 12 (3): e1005877.

Brunschwig, Hadassa, Liat Levi, Eyal Ben-David, Robert W. Williams, Benjamin Yakir, and Sagiv Shifman. 2012. “Fine-Scale Maps of Recombination Rates and Hotspots in the Mouse Genome.” Genetics 191 (3): 757–64.

Chan, Andrew H., Paul A. Jenkins, and Yun S. Song. 2012. “Genome-Wide Fine-Scale Recombination Rate Variation in Drosophila Melanogaster.” PLoS Genetics 8 (12): e1003090.

Chang, Christopher C., Carson C. Chow, Laurent Cam Tellier, Shashaank Vattikuti, Shaun M. Purcell, and James J. Lee. 2015. “Second-Generation PLINK: Rising to the Challenge of Larger and Richer Datasets.” GigaScience 4 (February): 7.

Cheung, Vivian G., Joshua T. Burdick, Deborah Hirschmann, and Michael Morley. 2007. “Polymorphic Variation in Human Meiotic Recombination.” American Journal of Human Genetics 80 (3): 526–30.

Chowdhury, Reshmi, Philippe R. J. Bois, Eleanor Feingold, Stephanie L. Sherman, and Vivian G. Cheung. 2009. “Genetic Analysis of Variation in Human Meiotic Recombination.” PLoS Genetics 5 (9): e1000648.

Cirulli, Elizabeth T., Richard M. Kliman, and Mohamed A. F. Noor. 2007. “Fine-Scale Crossover Rate Heterogeneity in Drosophila Pseudoobscura.” Journal of Molecular Evolution 64 (1): 129–35.

Cohen-Zinder, Miri, Eyal Seroussi, Denis M. Larkin, Juan J. Loor, Annelie Everts-van der Wind, Jun-Heon Lee, James K. Drackley, et al. 2005. “Identification of a Missense Mutation in the Bovine ABCG2 Gene with a Major Effect on the QTL on Chromosome 6 Affecting Milk Yield and Composition in Holstein Cattle.” Genome Research 15 (7): 936–44.

Coop, Graham, Xiaoquan Wen, Carole Ober, Jonathan K. Pritchard, and Molly Przeworski. 2008. “High-Resolution Mapping of Crossovers Reveals Extensive Variation in Fine-Scale Recombination Patterns among Humans.” Science 319 (5868): 1395–98.

Cox, Allison, Cheryl L. Ackert-Bicknell, Beth L. Dumont, Yueming Ding, Jordana Tzenova Bell, Gudrun A. Brockmann, Jon E. Wergedal, et al. 2009. “A New Standard Genetic Map for the Laboratory Mouse.” Genetics 182 (4): 1335–44.

Crawford, Dana C., Tushar Bhangale, Na Li, Garrett Hellenthal, Mark J. Rieder, Deborah A. Nickerson, and Matthew Stephens. 2004. “Evidence for Substantial Fine-Scale Variation in Recombination Rates across the Human Genome.” Nature Genetics 36 (7). Nature Publishing Group: 700–706.

Druet, Tom, and Michel Georges. 2015. “LINKPHASE3: An Improved Pedigree-Based Phasing Algorithm Robust to Genotyping and Map Errors.” Bioinformatics 31 (10): 1677–79.

Fariello, Maria-Ines, Bertrand Servin, Gwenola Tosser-Klopp, Rachel Rupp, Carole Moreno, International Sheep Genomics Consortium, Magali San Cristobal, and Simon Boitard. 2014. “Selection Signatures in Worldwide Sheep Populations.” PloS One 9 (8): e103813.

Ferrer-Admetlla, Anna, Mason Liang, Thorfinn Korneliussen, and Rasmus Nielsen. 2014. “On Detecting Incomplete Soft or Hard Selective Sweeps Using Haplotype Structure.” Molecular Biology and Evolution 31 (5): 1275–91.

Fraley, Chris, and Adrian E. Raftery. 2002. “Model-Based Clustering, Discriminant Analysis, and Density Estimation.” Journal of the American Statistical Association 97 (458). Taylor & Francis: 611–31.

Groenen, Martien A. M., Alan L. Archibald, Hirohide Uenishi, Christopher K. Tuggle, Yasuhiro Takeuchi, Max F. Rothschild, Claire Rogel-Gaillard, et al. 2012. “Analyses of Pig Genomes Provide Insight into Porcine Demography and Evolution.” Nature 491 (7424): 393–98.

Groenen, Martien A. M., Per Wahlberg, Mario Foglio, Hans H. Cheng, Hendrik-Jan Megens, Richard P. M. A. Crooijmans, Francois Besnier, et al. 2009. “A High-Density SNP-Based Linkage Map of the Chicken Genome Reveals Sequence Features Correlated with Recombination Rate.” Genome Research 19 (3): 510–19.

Guan, Yongtao, and Matthew Stephens. 2008. “Practical Issues in Imputation-Based Association Mapping.” PLoS Genetics 4 (12): e1000279.

Hassold, Terry, Heather Hall, and Patricia Hunt. 2007. “The Origin of Human Aneuploidy: Where We Have Been, Where We Are Going.” Human Molecular Genetics 16 (R2): R203–8.

Howie, Bryan N., Peter Donnelly, and Jonathan Marchini. 2009. “A Flexible and Accurate Genotype Imputation Method for the next Generation of Genome-Wide Association Studies.” PLoS Genetics 5 (6): e1000529.

Ignatieva, Elena V., Victor G. Levitsky, Nikolay S. Yudin, Mikhail P. Moshkin, and Nikolay A. Kolchanov. 2014. “Genetic Basis of Olfactory Cognition: Extremely High Level of DNA Sequence Polymorphism in Promoter Regions of the Human Olfactory Receptor Genes Revealed Using the 1000 Genomes Project Dataset.” Frontiers in Psychology 5 (March): 247.

International HapMap Consortium, Kelly A. Frazer, Dennis G. Ballinger, David R. Cox, David A. Hinds, Laura L. Stuve, Richard A. Gibbs, et al. 2007. “A Second Generation Human Haplotype Map of over 3.1 Million SNPs.” Nature 449 (7164): 851–61.

Jiang, Yu, Min Xie, Wenbin Chen, Richard Talbot, Jillian F. Maddox, Thomas Faraut, Chunhua Wu, et al. 2014. “The Sheep Genome Illuminates Biology of the Rumen and Lipid Metabolism.” Science 344 (6188): 1168–73.

Johnston, S. E., J. Huisman, P. A. Ellis, and J. M. Pemberton. 2017. “A High-Density Linkage Map Reveals Sexually-Dimorphic Recombination Landscapes in Red Deer (Cervus Elaphus).” bioRxiv. biorxiv.org. http://biorxiv.org/content/early/2017/01/13/100131.abstract.

Johnston, Susan E., Camillo Bérénos, Jon Slate, and Josephine M. Pemberton. 2016. “Conserved Genetic Architecture Underlying Individual Recombination Rate Variation in a Wild Population of Soay Sheep (Ovis Aries).” Genetics 203 (1): 583–98.

Johnston, Susan E., Jacob Gratten, Camillo Berenos, Jill G. Pilkington, Tim H. Clutton-Brock, Josephine M. Pemberton, and Jon Slate. 2013. “Life History Trade-Offs at a Single Locus Maintain Sexually Selected Genetic Variation.” Nature 502 (7469): 93–95.

Kadri, Naveen Kumar, Chad Harland, Pierre Faux, Nadine Cambisano, Latifa Karim, Wouter Coppieters, Sébastien Fritz, et al. 2016a. “Coding and Noncoding Variants in HFM1, MLH3, MSH4, MSH5, RNF212, and RNF212B Affect Recombination Rate in Cattle.” Genome Research 26 (10): 1323–32.

Kadri, Naveen Kumar, Chad Harland, Pierre Faux, Nadine Cambisano, Latifa Karim, Wouter Coppieters, Sébastien Fritz, et al. 2016b. “Coding and Noncoding Variants in HFM1, MLH3, MSH4, MSH5, RNF212, and RNF212B Affect Recombination Rate in Cattle.” Genome Research 26 (10): 1323–32.

Kari, Vijayalakshmi, Andrei Shchebet, Heinz Neumann, and Steven A. Johnsen. 2011. “The H2B Ubiquitin Ligase RNF40 Cooperates with SUPT16H to Induce Dynamic Changes in Chromatin Structure during DNA Double-Strand Break Repair.” Cell Cycle 10 (20): 3495–3504.

Kaur, Taniya, and Matthew V. Rockman. 2014. “Crossover Heterogeneity in the Absence of Hotspots in Caenorhabditis Elegans.” Genetics 196 (1): 137–48.

Kijas, James, J. Lenstra, B. Hayes, S. Boitard, Porto Neto L, Magali San Cristobal, Bertrand Servin, et al. 2012. “Genome-Wide Analysis of the World’s Sheep Breeds Reveals High Levels of Historic Mixture and Strong Recent Selection.” PLoS Biology. journals.plos.org. http://journals.plos.org/plosbiology/article?id=10.1371/journal.pbio.1001258.

Kim, E-S, A. R. Elbeltagy, A. M. Aboul-Naga, B. Rischkowsky, B. Sayre, J. M. Mwacharo, and M. F. Rothschild. 2016. “Multiple Genomic Signatures of Selection in Goats and Sheep Indigenous to a Hot Arid Environment.” Heredity 116 (3): 255–64.

Kong, A., G. Thorleifsson, H. Stefansson, G. Masson, A. Helgason, D. F. Gudbjartsson, G. M. Jonsdottir, et al. 2008a. “Sequence Variants in the RNF212 Gene Associate with Genome-Wide Recombination Rate.” Science 319 (5868): 1398–1401.

Kong, A., G. Thorleifsson, H. Stefansson, G. Masson, A. Helgason, D. F. Gudbjartsson, G. M. Jonsdottir, et al. 2008b. “Sequence Variants in the RNF212 Gene Associate with Genome-Wide Recombination Rate.” Science 319 (5868): 1398–1401.

Kong, Augustine, Gudmar Thorleifsson, Michael L. Frigge, Gisli Masson, Daniel F. Gudbjartsson, Rasmus Villemoes, Erna Magnusdottir, Stefania B. Olafsdottir, Unnur Thorsteinsdottir, and Kari Stefansson. 2014. “Common and Low-Frequency Variants Associated with Genome-Wide Recombination Rate.” Nature Genetics 46 (1): 11–16.

Kong, Augustine, Gudmar Thorleifsson, Daniel F. Gudbjartsson, Gisli Masson, Asgeir Sigurdsson, Aslaug Jonasdottir, G. Bragi Walters, et al. 2010. “Fine-Scale Recombination Rate Differences between Sexes, Populations and Individuals.” Nature 467 (7319): 1099–1103.

Lange, Julian, Shintaro Yamada, Sam E. Tischfield, Jing Pan, Seoyoung Kim, Xuan Zhu, Nicholas D. Socci, Maria Jasin, and Scott Keeney. 2016. “The Landscape of Mouse Meiotic Double-Strand Break Formation, Processing, and Repair.” Cell 167 (3): 695–708.e16.

Li, Heng, and Richard Durbin. 2011. “Inference of Human Population History from Individual Whole-Genome Sequences.” Nature 475 (7357): 493–96.

Li, Na, and Matthew Stephens. 2003. “Modeling Linkage Disequilibrium and Identifying Recombination Hotspots Using Single-Nucleotide Polymorphism Data.” Genetics 165 (4): 2213–33.

Ma, Li, Jeffrey R. O’Connell, Paul M. VanRaden, Botong Shen, Abinash Padhi, Chuanyu Sun, Derek M. Bickhart, et al. 2015. “Cattle Sex-Specific Recombination and Genetic Control from a Large Pedigree Analysis.” PLoS Genetics 11 (11): e1005387.

Mancera, Eugenio, Richard Bourgon, Alessandro Brozzi, Wolfgang Huber, and Lars M. Steinmetz. 2008. “High-Resolution Mapping of Meiotic Crossovers and Non-Crossovers in Yeast.” Nature 454 (7203): 479–85.

McVean, Gil, Philip Awadalla, and Paul Fearnhead. 2002. “A Coalescent-Based Method for Detecting and Estimating Recombination from Gene Sequences.” Genetics 160 (3): 1231–41.

Mihola, Ondrej, Zdenek Trachtulec, Cestmir Vlcek, John C. Schimenti, and Jiri Forejt. 2009. “A Mouse Speciation Gene Encodes a Meiotic Histone H3 Methyltransferase.” Science 323 (5912): 373–75.

Moreno-Romieux, Carole, Flavie Tortereau, Jérome Raoul, and Bertrand Servin. 2017. “High Density Genotypes of French Sheep Populations.” doi: 10.5281/zenodo.237116.

Myers, Simon, Leonardo Bottolo, Colin Freeman, Gil McVean, and Peter Donnelly. 2005. “A Fine-Scale Map of Recombination Rates and Hotspots across the Human Genome.” Science 310 (5746): 321–24.

Myers, S., C. C. A. Spencer, A. Auton, L. Bottolo, C. Freeman, P. Donnelly, and G. McVean. 2006. “The Distribution and Causes of Meiotic Recombination in the Human Genome.” Biochemical Society Transactions 34 (Pt 4): 526–30.

Nagamine, Yoshitaka, Ricardo Pong-Wong, Pau Navarro, Veronique Vitart, Caroline Hayward, Igor Rudan, Harry Campbell, et al. 2012. “Localising Loci Underlying Complex Trait Variation Using Regional Genomic Relationship Mapping.” PloS One 7 (10). Public Library of Science: e46501.

Nagel, Anja C., Patrick Fischer, Jutta Szawinski, Martina K. La Rosa, and Anette Preiss. 2012. “Cyclin G Is Involved in Meiotic Recombination Repair in Drosophila Melanogaster.” Journal of Cell Science 125 (Pt 22): 5555–63.

Norris, Belinda J., and Vicki A. Whan. 2008. “A Gene Duplication Affecting Expression of the Ovine ASIP Gene Is Responsible for White and Black Sheep.” Genome Research 18 (8): 1282–93.

O’Reilly, Paul F., Ewan Birney, and David J. Balding. 2008. “Confounding between Recombination and Selection, and the Ped/Pop Method for Detecting Selection.” Genome Research 18 (8): 1304–13.

Pratto, Florencia, Kevin Brick, Pavel Khil, Fatima Smagulova, Galina V. Petukhova, and R. Daniel Camerini-Otero. 2014. “Recombination Initiation Maps of Individual Human Genomes.” Science 346 (6211). American Association for the Advancement of Science: 1256442.

Qiao, Huanyu, H. B. D. Prasada Rao, Ye Yang, Jared H. Fong, Jeffrey M. Cloutier, Dekker C. Deacon, Kathryn E. Nagel, et al. 2014. “Antagonistic Roles of Ubiquitin Ligase HEI10 and SUMO Ligase RNF212 Regulate Meiotic Recombination.” Nature Genetics 46 (2): 194–99.

Rao, H. B. D. Prasada, Huanyu Qiao, Shubhang K. Bhatt, Logan R. J. Bailey, Hung D. Tran, Sarah L. Bourne, Wendy Qiu, et al. 2016. “A SUMO-Ubiquitin Relay Recruits Proteasomes to Chromosome Axes to Regulate Meiotic Recombination.” bioRxiv. Cold Spring Harbor Labs Journals. doi: 10.1101/095711.

Reynolds, April, Huanyu Qiao, Ye Yang, Jefferson K. Chen, Neil Jackson, Kajal Biswas, J. Kim Holloway, et al. 2013. “RNF212 Is a Dosage-Sensitive Regulator of Crossing-over during Mammalian Meiosis.” Nature Genetics 45 (3): 269–78.

Ritz, Kathryn R., Mohamed A. F. Noor, and Nadia D. Singh. 2017. “Variation in Recombination Rate: Adaptive or Not?” Trends in Genetics: TIG 33 (5): 364–74.

Rochus, Christina Marie, Flavie Tortereau, Florence Plisson-Petit, Gwendal Restoux, Carole Moreno-Romieux, Gwenola Tosser-Klopp, and Bertrand Servin. 2017. “High Density Genome Scan for Selection Signatures in French Sheep Reveals Allelic Heterogeneity and Introgression at Adaptive Loci.” bioRxiv. Cold Spring Harbor Labs Journals. doi: 10.1101/103010.

Rockman, Matthew V., and Leonid Kruglyak. 2009. “Recombinational Landscape and Population Genomics of Caenorhabditis Elegans.” PLoS Genetics 5 (3): e1000419.

Rosa, H. J. D., and M. J. Bryant. 2003. “Seasonality of Reproduction in Sheep.” Small Ruminant Research: The Journal of the International Goat Association 48 (3): 155–71.

Ruiz-Herrera, Aurora, Miluse Vozdova, Jonathan Fernández, Hana Sebestova, Laia Capilla, Jan Frohlich, Covadonga Vara, et al. 2017. “Recombination Correlates with Synaptonemal Complex Length and Chromatin Loop Size in Bovids-Insights into Mammalian Meiotic Chromosomal Organization.” Chromosoma, January. doi: 10.1007/s00412-016-0624-3.

Sabeti, Pardis C., David E. Reich, John M. Higgins, Haninah Z. P. Levine, Daniel J. Richter, Stephen F. Schaffner, Stacey B. Gabriel, et al. 2002. “Detecting Recent Positive Selection in the Human Genome from Haplotype Structure.” Nature 419 (6909): 832–37.

Sandor, Cynthia, Wanbo Li, Wouter Coppieters, Tom Druet, Carole Charlier, and Michel Georges. 2012a. “Genetic Variants in REC8, RNF212, and PRDM9 Influence Male Recombination in Cattle.” PLoS Genetics 8 (7): e1002854.

Sandor, Cynthia, Wanbo Li, Wouter Coppieters, Tom Druet, Carole Charlier, and Michel Georges. 2012b. “Genetic Variants in REC8, RNF212, and PRDM9 Influence Male Recombination in Cattle.” PLoS Genetics 8 (7): e1002854.

Scheet, Paul, and Matthew Stephens. 2006. “A Fast and Flexible Statistical Model for Large-Scale Population Genotype Data: Applications to Inferring Missing Genotypes and Haplotypic Phase.” American Journal of Human Genetics 78 (4): 629–44.

Servin, Bertrand, and Matthew Stephens. 2007. “Imputation-Based Analysis of Association Studies: Candidate Regions and Quantitative Traits.” PLoS Genetics 3 (7): e114.

Shifman, Sagiv, Jordana Tzenova Bell, Richard R. Copley, Martin S. Taylor, Robert W. Williams, Richard Mott, and Jonathan Flint. 2006. “A High-Resolution Single Nucleotide Polymorphism Genetic Map of the Mouse Genome.” PLoS Biology 4 (12): e395.

Słabicki, Mikołaj, Mirko Theis, Dragomir B. Krastev, Sergey Samsonov, Emeline Mundwiller, Magno Junqueira, Maciej Paszkowski-Rogacz, et al. 2010. “A Genome-Scale DNA Repair RNAi Screen Identifies SPG48 as a Novel Gene Associated with Hereditary Spastic Paraplegia.” PLoS Biology 8 (6): e1000408.

Stathopoulos, Sofia, Jacqueline M. Bishop, and Colleen O’Ryan. 2014. “Genetic Signatures for Enhanced Olfaction in the African Mole-Rats.” PloS One 9 (4): e93336.

Stephens, Matthew. 2017. “False Discovery Rates: A New Deal.” Biostatistics 18 (2): 275–94.

Stevison, Laurie S., and Mohamed A. F. Noor. 2010. “Genetic and Evolutionary Correlates of Fine-Scale Recombination Rate Variation in Drosophila Persimilis.” Journal of Molecular Evolution 71 (5-6): 332–45.

Storey, J. D., and R. Tibshirani. 2003. “Statistical Significance for Genomewide Studies.” Proceedings of the National Academy of Sciences 100 (16): 9440–45.

Sturtevant, A. H. 1913. “The Linear Arrangement of Six Sex-Linked Factors in Drosophila, as Shown by Their Mode of Association.” The Journal of Experimental Zoology 14 (1). Wiley Subscription Services, Inc., A Wiley Company: 43–59.

Takasuga, Akiko. 2015. “PLAG1 and NCAPG-LCORL in Livestock.” Animal Science Journal = Nihon Chikusan Gakkaiho. Wiley Online Library. http://onlinelibrary.wiley.com/doi/10.1111/asj.12417/pdf.

Tapanainen, J. S., K. Aittomäki, J. Min, T. Vaskivuo, and I. T. Huhtaniemi. 1997. “Men Homozygous for an Inactivating Mutation of the Follicle-Stimulating Hormone (FSH) Receptor Gene Present Variable Suppression of Spermatogenesis and Fertility.” Nature Genetics 15 (2): 205–6.

Tortereau, Flavie, Bertrand Servin, Laurent Frantz, Hendrik-Jan Megens, Denis Milan, Gary Rohrer, Ralph Wiedmann, et al. 2012. “A High Density Recombination Map of the Pig Reveals a Correlation between Sex-Specific Recombination and GC Content.” BMC Genomics 13 (November): 586.

Voight, Benjamin F., Sridhar Kudaravalli, Xiaoquan Wen, and Jonathan K. Pritchard. 2006. “A Map of Recent Positive Selection in the Human Genome.” PLoS Biology 4 (3): e72.

Wang, Jianbin, H. Christina Fan, Barry Behr, and Stephen R. Quake. 2012. “Genome-Wide Single-Cell Analysis of Recombination Activity and de Novo Mutation Rates in Human Sperm.” Cell 150 (2): 402–12.

Wang, Richard J., and Bret A. Payseur. 2017. “Genetics of Genome-Wide Recombination Rate Evolution in Mice from an Isolated Island.” Genetics, June. doi: 10.1534/genetics.117.202382.

Zeileis, Achim, and Gabor Grothendieck. 2005. “Zoo: S3 Infrastructure for Regular and Irregular Time Series.” Journal of Statistical Software, Articles 14 (6): 1–27.

Zhou, Xiang, Peter Carbonetto, and Matthew Stephens. 2013. “Polygenic Modeling with Bayesian Sparse Linear Mixed Models.” PLoS Genetics 9 (2): e1003264.

